# Cascade Diversification Directs the Generation of Neuronal Diversity in Hypothalamus

**DOI:** 10.1101/2020.06.01.125054

**Authors:** Yu-Hong Zhang, Mingrui Xu, Si Li, Haoda Wu, Xiang Shi, Xize Guo, Wenhui Mu, Ling Gong, Mingze Yao, Miao He, Qing-Feng Wu

**Affiliations:** State Key Laboratory of Molecular Development Biology, Institute of Genetics and Developmental Biology, Chinese Academy of Sciences, Beijing, 100101, China; University of Chinese Academy of Sciences, Beijing 100101, China; Institutes of Brain Science, State Key Laboratory of Medical Neurobiology and MOE Frontiers Center for Brain Science, Fudan University, Shanghai 200032, China; Institutes of Biomedical Sciences, Shanxi University, Taiyuan 030006, China.; CAS Center for Excellence in Brain Science and Intelligence Technology, Shanghai 200031, China

## Abstract

The hypothalamus contains an astounding heterogeneity of neurons to achieve its role in regulating endocrine, autonomic and behavioral functions. Despite previous progress in deciphering the gene regulatory programs linked to hypothalamus development, its molecular developmental trajectory and origin of neuronal diversity remain largely unknown. Here we combine transcriptomic profiling of 43,261 cells derived from Rax^+^ hypothalamic neuroepithelium with lineage tracing to map a developmental landscape of mouse hypothalamus and delineate the developmental trajectory of radial glial cells (RGCs), intermediate progenitor cells (IPCs), nascent neurons and peptidergic neurons in the lineage hierarchy. We show that RGCs adopt a conserved strategy for multipotential differentiation but generate both Ascl1^+^ and Neurog2^+^ IPCs, which display regionally differential origins in telencephalon. As transit-amplifying cells, Ascl1^+^ IPCs differ from their telencephalic counterpart by displaying fate bifurcation to produce both glutamatergic and GABAergic neurons. After classifying the developing neurons into 29 subtypes coded by diverse transcription factors, neurotransmitters and neuropeptides, we identified their molecular determinants via regulon analysis and further found that postmitotic neurons at nascent state possess the potential to resolve into more diverse subtypes of peptidergic neurons. Together, our study offers a single-cell framework for hypothalamus development and reveals that multiple cell types along the order of lineage hierarchy contribute to the fate diversification of hypothalamic neurons in a stepwise fashion, suggesting that a cascade diversifying model can deconstruct the origin of neuronal diversity.

## INTRODUCTION

A mechanistic understanding of brain development requires a systematic survey of neural progenitor cell types, their lineage specification and developmental maturation of postmitotic neurons. Cumulative evidences based on single-cell transcriptomic analysis have revealed the transcriptional heterogeneity of cortical neural progenitors, their temporal patterning and the differentiation trajectories of both excitatory neurons and inhibitory interneurons in the developing mammalian neocortex (Johnson et al., 2015; Loo et al., 2019; Mayer et al., 2018; Mi et al., 2018; Nowakowski et al., 2017; Zhong et al., 2018). However, the developmental hierarchy of hypothalamus, representing a conserved but extremely diverse and complex brain structure, remains poorly understood.

Hypothalamus maintains systemic homeostasis of animals by regulating endocrine, autonomic and behavioral functions, ranging from hunger, sleep, thirst, circadian rhythm and body temperature to mood regulation, sex drive and hormonal release (Puelles, 2012; Sternson, 2013). The functional complexity of hypothalamus relies on the extreme neuronal diversity generated during brain development. Hypothalamic neurons are comprised of magnocellular endocrine neurons (OXT, AVP, etc.), parvocellular secretory neurons (TRH, CRH, etc.), large peptidergic projection neurons (HCRT, MCH, etc.), parvocellular peptidergic neurons (POMC, AgRP, etc.) and other inhibitory or excitatory local-circuit neurons (Romanov et al., 2019). Various neuropeptides have long served as a basis for defining neuronal subtypes and understanding hypothalamic functionality. In contrast to numerous studies revealing the circuit and function of diverse hypothalamic neurons, little is known about how neuronal subtype-specific heterogeneity emerges from common radial glial cells (RGCs), acting as embryonic neural stem cells during development. There have been two models proposed for the origin and process of generating neuronal diversity in the mammalian brain. One model proposes that cortical RGCs sequentially produce fate-predetermined intermediate progenitor cells (IPCs) differentiating into deep-layer and upper-layer excitatory neurons (Kohwi and Doe, 2013). Recent single-cell analyses further support that the diversity of cortical inhibitory interneurons is already predetermined at the level of progenitors or shortly after becoming postmitotic (Mayer et al., 2018; Mi et al., 2018). Notably, the origins of excitatory glutamatergic and inhibitory GABAergic neurons, as well as their immediate progenitors, are spatially segregated and diverse in the neocortex (Marin and Muller, 2014). The second stochastic model postulates that retinal progenitor cells stochastically adopt diverse cell fate during differentiation (Gomes et al., 2011; He et al., 2012). Given the intermingling state of molecularly diverse neurons and their critical roles in maintaining homeostatic control, it is fundamentally important to investigate the developmental diversification and trajectories of hypothalamic neurons.

Here we sorted hypothalamic cells arising from Rax lineage for single-cell RNA sequencing and delineated the developmental trajectory of RGCs, IPCs, nascent neurons and peptidergic neurons within the neural lineage hierarchy. In contrast to the fate-predetermined model in cortex and stochastic model in retina, our transcriptomic analyses in combination with lineage tracing data support a cascade diversifying model wherein RGCs, IPCs and nascent neurons prompt the diversification of hypothalamic neuronal fate in a stepwise manner, uncovering a novel strategy for neural progenitors to generate an extreme neuronal diversity. We also identified the embryonic origin of postnatal tanycytes and the regulons specifying neuronal subtypes, which further provides a developmental perspective to understand hypothalamus plasticity and obtain valuable insights into hypothalamic disease such as anorexia, narcolepsy and insomnia.

## RESULTS

### Transcriptional Profiling of Developing Hypothalamus

Rax-CreER^T2^ tool mice have recently been used to label hypothalamic primordium at early embryonic stage (Pak et al., 2014). To infer the lineage progression underlying hypothalamus development, we first labeled hypothalamic neuroepithelium by applying a single dose of tamoxifen in *Rax-CreER^T2^::Ai14* mice at embryonic day 9 (E9) and performed lineage tracing for cell collection (Figure 1A and S1A). Next, we microdissected hypothalamus from induced *Rax-CreER^T2^::Ai14* mice across 4 different time points that cover the process of neurogenesis and early maturation for hypothalamic neurons [E11, E14, postnatal days 0 (P0) and P7], and collected live tdTomato^+^ cells by fluorescent-activated cell sorting for single-cell transcriptomic sequencing (Figures S1B-E). A final dataset of 43261 Rax-derived hypothalamic cells, covering an average of 11376 unique molecular identifiers (UMIs) and 3753 genes per cell, was obtained after quality control and batch integration (Figures S1F-M).

**Figure 1.**
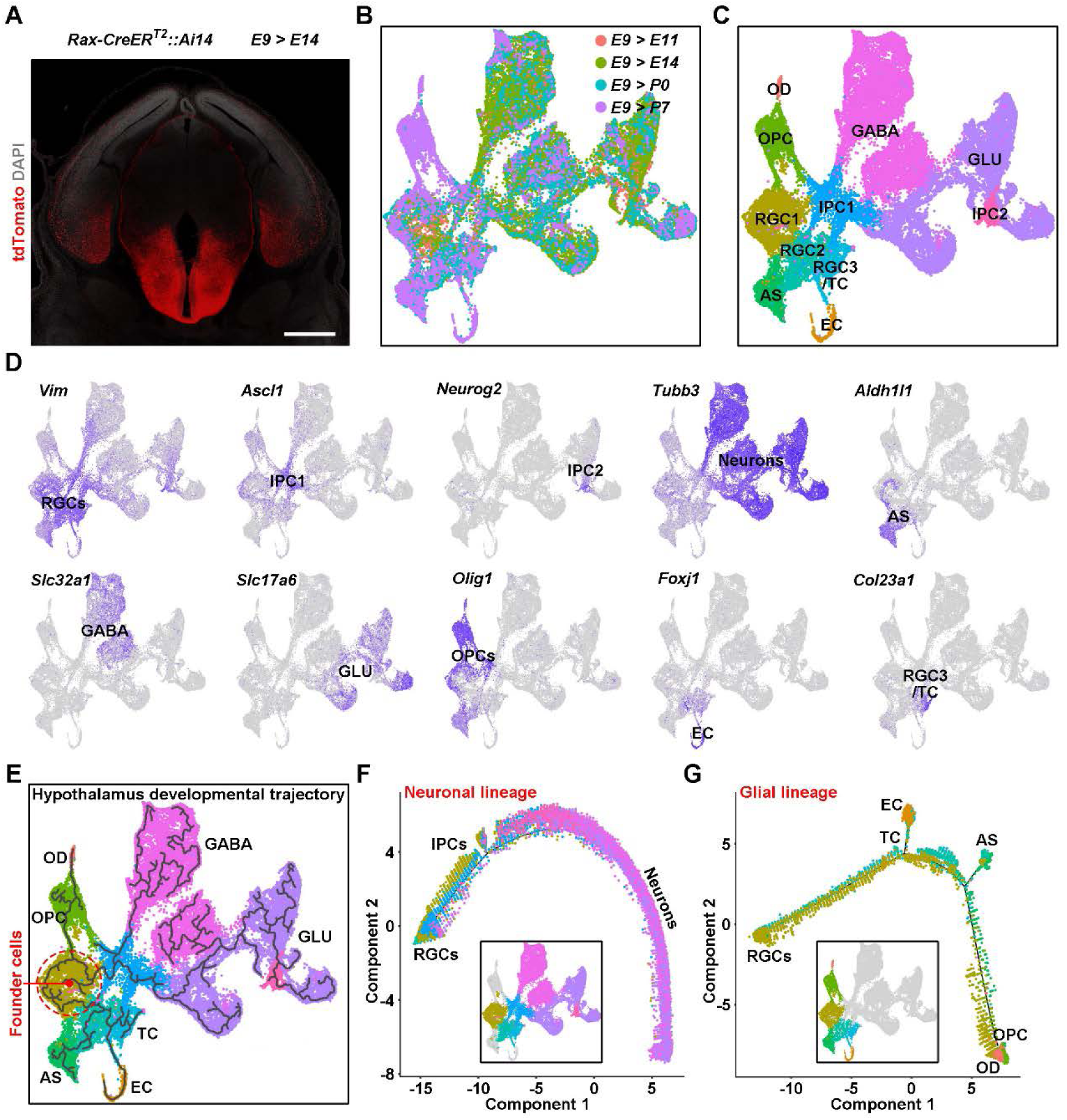
Molecular Developmental Trajectory of Mouse Hypothalamus. (A) Representative image showing the genetically-mediated fluorescent labeling of hypothalamic cell lineage with *Rax-CreER^T2^::Ai14* mice. The mice were induced with tamoxifen at a single dose of 132 mg/kg body weight at E9 and collected at E14. Scale bar, 500 µm. (B-C) UMAP visualization of 43,261 hypothalamic cells derived from Rax^+^ neuroepithelium, colored by the age (B) and groups (C). The hypothalamic cells were induced at E9 in *Rax-CreER^T2^::Ai14* mice and collected at E11, E14, P0 and P7 for fluorescence-activated cell sorting of tdTomato^+^ cells. The identity of cell groups are annotated according to known cell types. RGC, radial glial cells; IPC, intermediate progenitor cells; TC, tanycytes; EC, ependymal cells; AS, astrocytes; OPC, oligodendrocyte precursor cells; OD, oligodendrocytes; GABA, GABAergic neruons; GLU, glutamatergic neurons. (D) Imputed gene expression of canonical markers overlaid on UMAP plot. (E) Global developmental architecture of hypothalamic cell lineages inferred by Monocle 3. Red dashed circle indicates the potential founder cells giving rise to hypothalamus and black solid lines suggest the direction of lineage progression of hypothalamic cells. (F-G) Pseudotemporal trajectories of neuronal lineage (F) and glial lineage (G) in hypothalamus. The colored cells within boxed UMAP plots were used for imputing developmental trajectories.

To determine the major cell types in the developing hypothalamus, we employed feature selection, principal component analysis (PCA) and uniform manifold approximation and projection (UMAP) method for dimensionality reduction to cluster cells based on their transcriptomic similarity. UMAP stratified cells by their developmental timing and identities (Figures 1B, 1C and S2A). Mapping well-established marker genes onto the UMAP representation identified 8 principal cell types and 11 cell clusters: RGCs (*Vim^+^*), IPCs (*Ascl1^+^* or *Neurog2^+^*), glutamatergic neurons (GLU, *Tubb3^+^Slc17a6^+^*), GABAergic neurons (GABA, *Tubb3^+^Slc32a1^+^*), astrocytes (AS, *Aldh1l1^+^*), oligodendrocyte precursor cells (OPCs, *Olig1^+^Pdgfra^+^*), oligodendrocytes (OD, *Mbp^+^*) and ependymal cells (ECs, *Foxj1^+^*) (Figures 1C, 1D and S2B). Notably, a cluster of RGCs encompassed *Col23a1*^+^ tanycytes (TCs) born at postnatal stage (Altman and Bayer, 1978; Chen et al., 2017), implying its predetermined fate for TC lineage. The identity of cell clusters was further confirmed by a set of cell-type specific genes ascertained by differential gene expression analysis (Figure S2C; Table S1). Overall, the cell atlas captures cellular heterogeneity within the developing hypothalamus, and assigning each cluster to an established cell type allows us to analyze the temporal progression of neuronal and glial lineages in the hypothalamus.

As expected, prenatal cells were mainly featured by the process of cell division, pattern specification, neuron differentiation, axonogenesis and axon guidance, while postnatal cells were characterized by neuronal apoptosis, gliogenesis, myelination, synapse organization and maturation (Figure S2D; Table S2). The temporal analysis of cell-type proportion also corroborated the general rule that neurogenic processes preceded gliogenesis (Figure S2E). We then performed pseudotemporal analysis to reconstruct the developmental trajectory of hypothalamus across the entire dataset. Indeed, the multibranched cellular trajectory on the UMAP representation showed the multipotential differentiation of founder cells into GABAergic neurons, glutamatergic neurons, OPCs, AS, TCs and ECs (Figure 1E). We further subdivided all cells into neuronal and glial lineages for trajectory inference. The neuronal lineage analysis uncovered a simple continuous trajectory connecting RGCs, IPCs and neurons, whereas the pseudotemporal ordering of glial lineage cells indicated the multifurcation of RGCs into three branches: OPC-OD, AS and TC-EC sublineages (Figures 1F and 1G). Taken together, we combined cluster and trajectory analyses to stratify different neural cell types and unravel the developmental dynamics of hypothalamic cells.

### Temporal Dynamic Analysis of RGCs Reveals Embryonic Origin of Postnatal Tanycytes

To dissect the diversity among prenatal RGCs and postnatal RGC-like cells in hypothalamus, we subsetted all RGC clusters across different time points, regressed out cell cycle scores and identified 11 subclusters of cells (Figures 2A and S3A-D; Table S3 and S4). Using the expression pattern of lineage- and cell type-specific marker genes, we assigned a single identity to each cluster and subdivided the RGCs into two major subpopulations: RGCs in cycling, quiescent or transitional state without lineage preference (cRGC, qRGC or tRGC), and RGCs entering the primitive state of lineage differentiation toward IPCs, OPCs, astrocyte precursor (APs), ECs or TCs (pri-IPC, pri-OPC, pri-AP, pri-EC or pri-TC) (Figures 2A, S3E and S3F; Table S4). The trajectory inference further identified cRGCs as the major founder cells to transform into RGCs with lineage preference for IPC, OPC, AP and TC-EC (Figure 2B), suggesting that the fate of RGCs is predetermined before differentiation.

**Figure 2.**
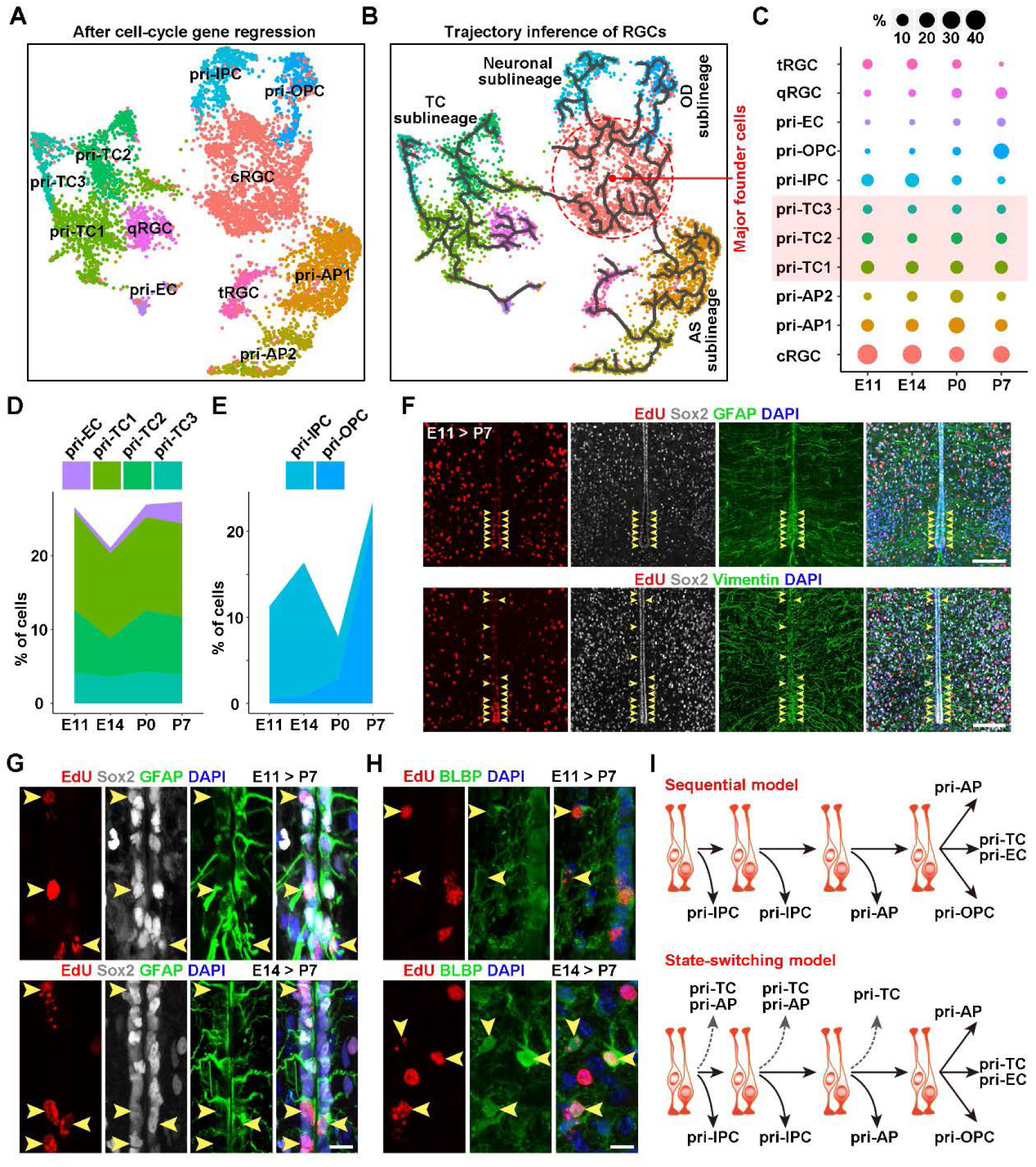
Strategically Conserved RGCs Switch into Quiescence as A Reservoir for Hypothalamic Tanycytes (A) UMAP visualization of 11 clusters of hypothalamic RGCs after cell-cycle gene regression, colored by cell subtypes and annotated based on the expression of putative markers for cycling, quiescence, IPCs, OPCs, TCs, ECs and astrocytes. cRGC, cycling RGCs; qRGC, quiescent RGCs; tRGC, transitional RGCs; pri-IPC, primitive IPCs; pri-OPC, primitive OPCs; pri-TC, primitive TCs; pri-EC, primitive ECs; pri-AP, primitive astrocyte precursors. Notably, cycling RGCs were still clustered together after regressing out cell cycle-related genes. (B) Pseudotemporal trajectory of RGCs overlaid on UMAP representation. The branched trajectory shows the differentiation of founder cells into neuronal, astroglial, oligodendrocyte and tanycyte sublineages. Red dashed circle indicates the starter cells. (C) The proportion of each RGC subtype at four developmental time points. The fractions of three pri-TC subtypes are highlighted in light red. (D-E) Temporal dynamics of the proportion of RGC subtypes committed to tanycyte (D) or neuronal and oligodendrocyte sublineages (E). (F) Sample confocal images showing the EdU^+^ label-retaining cells co-stained for tanycyte markers Sox2, GFAP and Vimentin along the third ventricle. The animals were pulsed with a single dose of EdU at E11 and sacrificed at P7 for brain tissue collection. Yellow arrowheads indicate the EdU^+^Sox2^+^GFAP^+^ or EdU^+^Sox2^+^Vimentin^+^ triple-labeled cells. Scale bar, 100 µm. (G) Magnified images showing the EdU^+^ label-retaining TCs pulsed at E11 (top) or E14 (bottom) and chased at P7. Yellow arrowheads mark the triple-labeled cells lining the third ventricle. Scale bars, 10 µm. (H) Representative images stained for EdU and BLBP in P7 hypothalamus from mice receiving a single EdU pulse labeling at E11 (top) or E14 (bottom). The EdU^+^BLBP^+^ label-retaining APs are indicated by arrowheads. Scale bar, 10 µm. (I) Schematic diagrams for traditional “sequential” model and our “state-switching” model. In the sequential model, RGCs sequentially produce neurons at embryonic stages, glia during early postnatal stage and eventually hypothalamic TCs. Our state-switching model suggests that a subpopulation of RGCs stop dividing and switch into a primitive state of TCs, even APs, at early embryonic stage in the hypothalamus.

Hypothalamic TCs, a specialized type of ependymal cells, have recently been considered as postnatal somatic stem cells in the adult mammalian brain (Kokoeva et al., 2005; Lee et al., 2012). Although “set-aside” and “continuous” models have been proposed to interpret the identity of precursors to adult neural stem cells in lateral ventricles and hippocampus (Berg et al., 2019; Fuentealba et al., 2015), the embryonic origin of hypothalamic TCs remains unclear. Our temporal analysis of cell-type proportion showed that the percentage of various pri-TCs did not increase postnatally but remained constant during the developmental continuum (Figures 2C and 2D), implicating that a subpopulation of RGCs starts to switch into the precursors of TCs at early developmental stages. In contrast, the temporal dynamics of pri-IPC and pri-OPC proportions indicated a predominantly prenatal neurogenesis and a postnatal surge in oligodendrocytosis (Figure 2E). To validate the early state-switch from naive RGCs to pri-TCs, we applied a single dose of ethynyldeoxyuridine (EdU) in E11 or E14 mice and performed label retention assay at P7. Indeed, the birth dating analysis showed that a subset of RGCs exited cell cycle and switched into the TC fate, marked by Sox2/Vimentin/GFAP and positioned along the third ventricle, as early as E11 (Figures 2F and 2G). Likewise, a subpopulation of BLBP^+^ APs in hypothalamic parenchyma were born before E14 (Figure 2H). Therefore, our results support a “state-switching” model wherein a substantial fraction of TCs is derived from early embryonic RGCs in parallel to prenatal neurogenesis, rather than the “sequential” model in which RGCs sequentially produce neurons, glia and terminally differentiate into TCs (Figure 2I).

### Hypothalamic RGCs Give Rise to Differential IPC Subpopulations

IPCs act as secondary, transient amplifying neural progenitor cells, enabling rapid neurogenesis in the developing brain (Kowalczyk et al., 2009). In telencephalon, neurogenic IPCs producing excitatory and inhibitory neurons arise from RGCs residing in dorsal and ventral pallium, respectively (Marin and Muller, 2014). To investigate the potential of hypothalamic RGCs in immediate progeny production, we extracted the cells within neuronal lineage at early developmental stage to conduct pseudotemporal analyses using Monocle and RNA velocity. The developmental trajectory showed that RGCs bifurcated into two IPC sublineages for generating neurons within hypothalamic niche (Figure S4), suggesting that the starter cells initiate a lineage program to drive neurogenesis through two distinct types of IPCs.

### Two Subpopulations of IPCs with Distinct Molecular Profiles, Lineage Potentials and Spatial Distribution

Proneural factors Ascl1 and Neurog2 have previously been found to promote neuronal differentiation of progenitor cells and induce distinct neuronal subtype identities, specifying the fate of GABAergic and glutamatergic neurons in the telencephalon, respectively (Aydin et al., 2019; Fode et al., 2000; Parras et al., 2002). Here we identified two subpopulations of hypothalamic IPCs (HIPCs), respectively defined by Ascl1 and Neurog2 (Figures 1C and 1D), and revealed the coexistence of both Ascl1^+^ and Neurog2^+^ progenitor cells in the developing hypothalamus (Figure 3A). After subsetting all HIPCs across different time points and regressing out cell cycle variation, we reclustered the cells and further compared the molecular profiles between the two main groups of HIPCs (Figures 3B, 3C and S5A-C; Table S5). Our data showed that HIPC1 and HIPC2 were featured by distinct sets of transcription factors (TFs): Ascl1, Sox3, Gsx1 and Isl1 cooperated to define the identity of HIPC1, while Neurog2 orchestrated with Nhlh2, Neurod1 and Otp to govern HIPC2 identity (Figure 3C). Moreover, we found that HIPC1 and HIPC2 shared the enriched gene sets involved in neuronal fate specification, whereas they displayed differentially enriched cellular components even before differentiating into neurons, with HIPC1 expressing genes related to synapse formation for neuronal input and HIPC2 expressing genes associated with axonal growth for neuronal output (Figure S5D).

**Figure 3.**
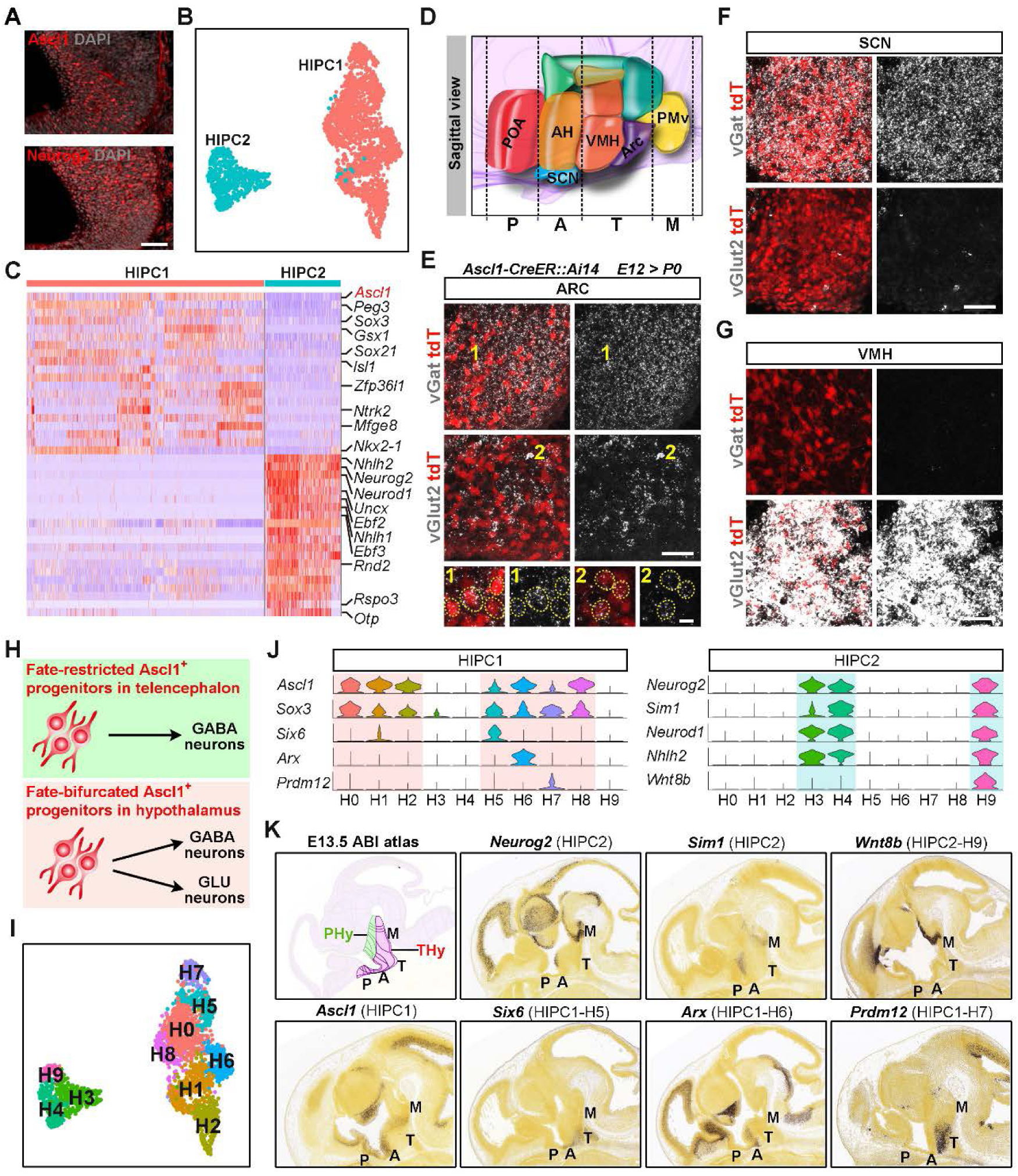
Fate Diversification of Two Subpopulations of IPCs (A) Representative images showing the co-existence of both Ascl1^+^ and Neurog2^+^ HIPCs in the E11 hypothalamus. Scale bar, 50 µm. (B) UMAP plot of two main subpopulations of hypothalamic IPCs (HIPCs). (C) Shown is the heatmap of genes differentially expressed in Ascl1^+^ HIPC1 and Neurog2^+^ HIPC2 groups. (D) Schematic diagraph showing the sagittal view of developing hypothalamus traversing preoptic (P), anterior (A), tuberal (T) and mammillary (M) zones. POA, preoptic area; AH, anterior hypothalamus; SCN, suprachiasmatic nucleus; VMH, ventromedial nucleus; ARC, arcuate nucleus; PMv, ventral premammilary nucleus. (E) The combination of single-molecule fluorescent *in situ* hybridization with immunostaining shows that Ascl1^+^ HIPC1 give rise to both vGat^+^ GABAergic and vGlut2^+^ glutamatergic neuronal progeny in ARC nucleus. *Ascl1-CreER^T2^::Ai14* mice were induced with tamoxifen at E12 for lineage tracing and sacrificed at P0 for analysis. The numbered, magnified images represent the colocalization of tdTomato with vGat or vGlut2 at single-cell resolution. Scale bars, 50 µm, and 10µm for magnified images. (F-G) Sample images showing the generation of pure vGat^+^ neurons in SCN (E) and vGlut2^+^ neurons in VMH (F) by Ascl1^+^ HIPC1 subtype. Scale bars, 50 µm. (H) Models depicting the fate restriction of Ascl1^+^ IPCs in telencephalon and the fate bifurcation of Ascl1^+^ IPCs in hypothalamus. (I) Subclustering of two main subpopulations of HIPCs. (J) Violin plots showing the representative genes specifically expressed in HIPC1, HIPC2 or their subtypes. Seven HIPC1 and three HIPC2 subtypes are highlighted with light red and cyan colors, respectively. (K) Spatial gene expression shown by *in situ* hybridization of sagittal brain sections from E13.5 mouse brains indicates the differential spatial distribution between HIPC1 and HIPC2. PHy, peduncular hypothalamus; THy, terminal hypothalamus. ABI atlas, Allen brain atlas.

While UMAP shows that HIPC2 was restricted within glutamatergic neuronal sublineage (Figure 1C, Figure S4B-C), the contiguity of HIPC1 with multiple neuronal sublineages raised the question related to the fate potentials of Ascl1^+^ IPCs in hypothalamus. Hence, we conducted lineage tracing using *Ascl1-CreER^T2^::Ai14* mice with tamoxifen induction at E12 to examine the fate potentials of HIPC1 (Figure S5E). Single-molecule RNA fluorescent *in situ* hybridization (smFISH) analysis revealed that Ascl1^+^ HIPCs contributed to the generation of both GABAergic (marked by vGat) and glutamatergic (marked by vGlut2) neuronal subtypes in multiple hypothalamic nuclei (Figures 3D-G and S5F-I), suggesting that Ascl1^+^ HIPCs are fate-bifurcated rather than fate-restricted in neuronal differentiation (Figure 3H).

We next sought to investigate the spatial distribution of distinct HIPC subpopulations in the embryonic hypothalamus. Reclustering analysis at higher resolution yielded 7 subclusters (H0-2 and H5-8) for HIPC1 and 3 subclusters (H3-4 and H9) for HIPC2, and UMAP-based trajectory inference implicates that the IPCs in H0-2 and H8 subgroups represented a more primitive state than the cells in H5-7 subclusters, confirmed by the expression analysis of stemness genes (Sparc, Slc1a3 and Apoe) (Figures 3I, S6A-D; Table S6). To explore the different spatial code of HIPC1 and HIPC2, we cross-referenced the markers with *in situ* hybridization data from the Allen Developing Mouse Brain Atlas (Figures 3J, 3K and S6E). The hypothalamus has traditionally been subdivided into peduncular and terminal regions along longitudinal axis, as well as preoptic, anterior, tuberal and mammillary zones along sagittal axis. Here we found that Neurog2^+^Sim1^+^ HIPC2 was spatially restricted within peduncular region and mammillary zone, whereas Ascl1^+^Sox3^+^ HIPC1 exhibited much more widespread distribution in preoptic, anterior and tuberal zones (Figure 3K). In particular, H5-7 subclusters representing a predifferentiated state spanned from anterior to tuberal zones (Figures 3K and S6C). Together, our results revealed the different molecular and spatial codes of two subpopulations of HIPCs, which may instruct their diversified lineage potential in hypothalamus.

### Hypothalamic Neuronal Subtypes and Their Regulons for Specifying Subtype Identities

To taxonomize the neuronal subtypes in the developing hypothalamus, we extracted all postmitotic neurons and subdivided them into two major neuronal classes: 13349 glutamatergic neurons and 13658 GABAergic neurons (Figure 4A). In the neocortex, Dlx homeobox TFs are required for GABAergic interneuron production and, in contrast, another four TFs (Fezf2, Ctip2, Tbr1 and Satb2) regulate the cell fate acquisition of glutamatergic cortical neurons (Leone et al., 2008; Petryniak et al., 2007). Notably, while Dlx factors including Dlx1, Dlx2, Dlx5 and Dlx6 were also enriched in differentiated GABAergic neurons, another set of TFs such as Nhlh2, Uncx, Fezf1 and Lhx5 showed class-specific expression in glutamatergic neurons within developing hypothalamus (Figure 4B), suggesting that cortical and hypothalamic neurons could adopt a comparable TF code to specify inhibitory neurons but different TF codes to drive the differentiation of excitatory neurons.

**Figure 4.**
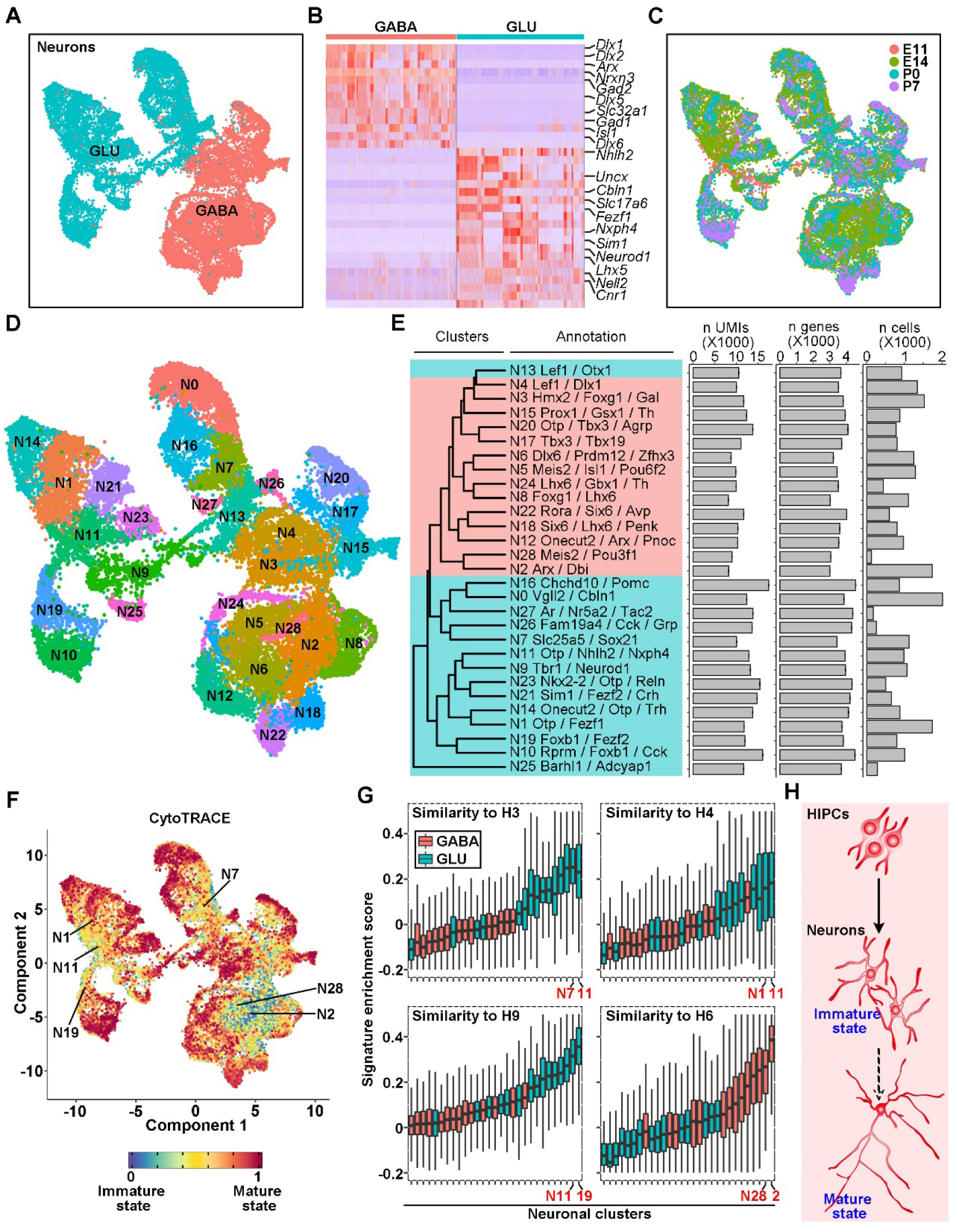
Subtypes and States of Postmitotic Neurons in the Developing Hypothalamus. (A) UMAP plot showing the two main groups of hypothalamic neurons distinguished by fast neurotransmitters. (B) Heatmap of genes differentially expressed in GABAergic and glutamatergic neurons in the developing hypothalalmus. (C) UMAP representation of postmitotic neurons colored by their developmental stages. (D) Subclustering of neurons into 29 subtypes, including 14 GABAergic and 15 glutamatergic subgroups. (E) Characterization and annotation of neuronal subtypes. Dendrogram shows the transcriptional relatedness of neuronal subtypes annotated by combinatorial molecular codes of transcription factors (TFs) and/or neuropeptides. The numbers of unique molecular identifiers (UMIs), detected genes and cells per cluster are shown in histograms (from left to right). Values of UMIs and genes represent mean ± SEM. (F) Maturation state of developing hypothalamic neurons predicted by CytoTRACE and overlaid on UMAP representation. Six clusters of neurons at relatively nascent state are labeled. (G) Box plots showing the gene set enrichment scores of HIPC molecular signatures for each neuronal subtypes. Boxes represent interquartile range and whiskers represent the minimum and maximum values. The top 2 neuronal subtypes with highest scores, displaying a transcriptional similarity to 4 HIPC subgroups (H3, H4, H9 and H6), are labeled for each box plot. (H) Schematic diagram shows that the differentiation of hypothalamic neurons requires a transition from nascent to mature state.

We subsequently analyzed the spatiotemporal organization of hypothalamic neurons. Along the temporal trajectories, the transcriptional programs of both neuronal classes changed from neuronal differentiation and pattern specification to synapse organization, neurotransmitter release and hormone secretion (Figures 4C and S7). We further catalogued 29 neuronal subtypes by unsupervised clustering approach, used a combinatorial code of TFs and neuropeptides to define the identity of each cluster, and linked the molecularly defined cell subtypes to their spatial distribution in diverse hypothalamic nuclei (Figures 4D, 4E, S8 and S9; Table S7). For example, neuronal clusters N3, N8, N24 and N28 were spatially segregated in preoptic area (POA, Figure S8A).

Moreover, we applied CytoTRACE to decipher the degree of differentiation of all neuronal subtypes (Gulati et al., 2020) and found a “central-to-peripheral” maturation pattern in both GABAergic and glutamatergic neurons on the UMAP representation (Figures 4F, S10A and S10B). The scoring of 29 neuronal subtypes with the molecular signatures of all HIPC subgroups indicated that glutamatergic neuronal clusters bore closer resemblance to HIPC2 subgroups (H3-4 and H9) and the neuronal subtypes with higher scores tended to be immature cells (Figures 4G and FS10C). These results suggest that our dataset covers a continuum of neurons in both immature and mature states (Figure 4H). Collectively, we provide a comprehensive atlas of developing hypothalamic neurons with different molecular profiles, spatial distribution and maturation degree.

We proceeded to explore the subtype-specific TFs and their essential roles in organizing the gene regulatory networks (i.e. regulons) that determine the fate of each neuronal subtype. Firstly, we screened 4482 transcriptional regulators and identified 191 TFs with specific expression in different neuronal subtypes (Figure 5A; Table S8). It has previously been reported that core sets of TFs act together in a regulated manner to control gene expression and shape cellular identities (Chen et al., 2008; Hobert, 2004; Shirasaki and Pfaff, 2002). We thereby performed co-expression modular analysis and found 15 TF modules which may function synergistically to define neuronal subtype identities (Figure 5B). We exemplified the TF module 11 wherein Lmx1a, Irx5, Pitx2, Barhl1 and Foxa1 displayed a significant association. A recent report demonstrated that Lmx1a, Pitx2 and Barhl1 are co-expressed in most neurons within subthalamic nucleus (STN) and exhibit permutations of co-expression in other hypothalamic glutamatergic neurons (Kee et al., 2017). Our smFISH analysis in conjunction with immunostaining validated the co-labeling of Lmx1a and Irx5 in multiple nuclei and confirmed the co-expression of Barhl1 and Pitx2 in STN (Figure 5C). Lastly, we identified 313 regulons with strikingly enriched motifs for the master TFs by applying cis-regulatory analysis to our single-cell dataset (Figure 5D), allowing us to determine the critical regulators for cell identity. We then assigned regulons to each of neuronal subtypes, ordered the regulons based on their regulon specificity score (RSS) and chose the top-ranked regulons for further analysis (Figures 5E-H and S11). To test the efficacy of this approach, we started with the neuronal subtypes (N22 and N18) distributed in suprachiasmatic nucleus (SCN) wherein the core gene regulatory network has been well studied (Clark et al., 2013; Sato et al., 2004; Wen et al., 2020). Indeed, our data revealed Rora (a critical regulator of clock genes) and Six6 (a master regulator of neuronal fate in SCN) as the active regulons associated with N22 and N18 neuronal subtypes, respectively (Figure 5E; Figure S11A). Further analysis showed the specific expression of top-ranked regulons in corresponding neuronal clusters (Figures 5E-H, Figure S11A-J). Taken together, we provide a taxonomy of developing hypothalamic neurons at nascent or mature state and a list of essential regulons specifying their identity.

**Figure 5.**
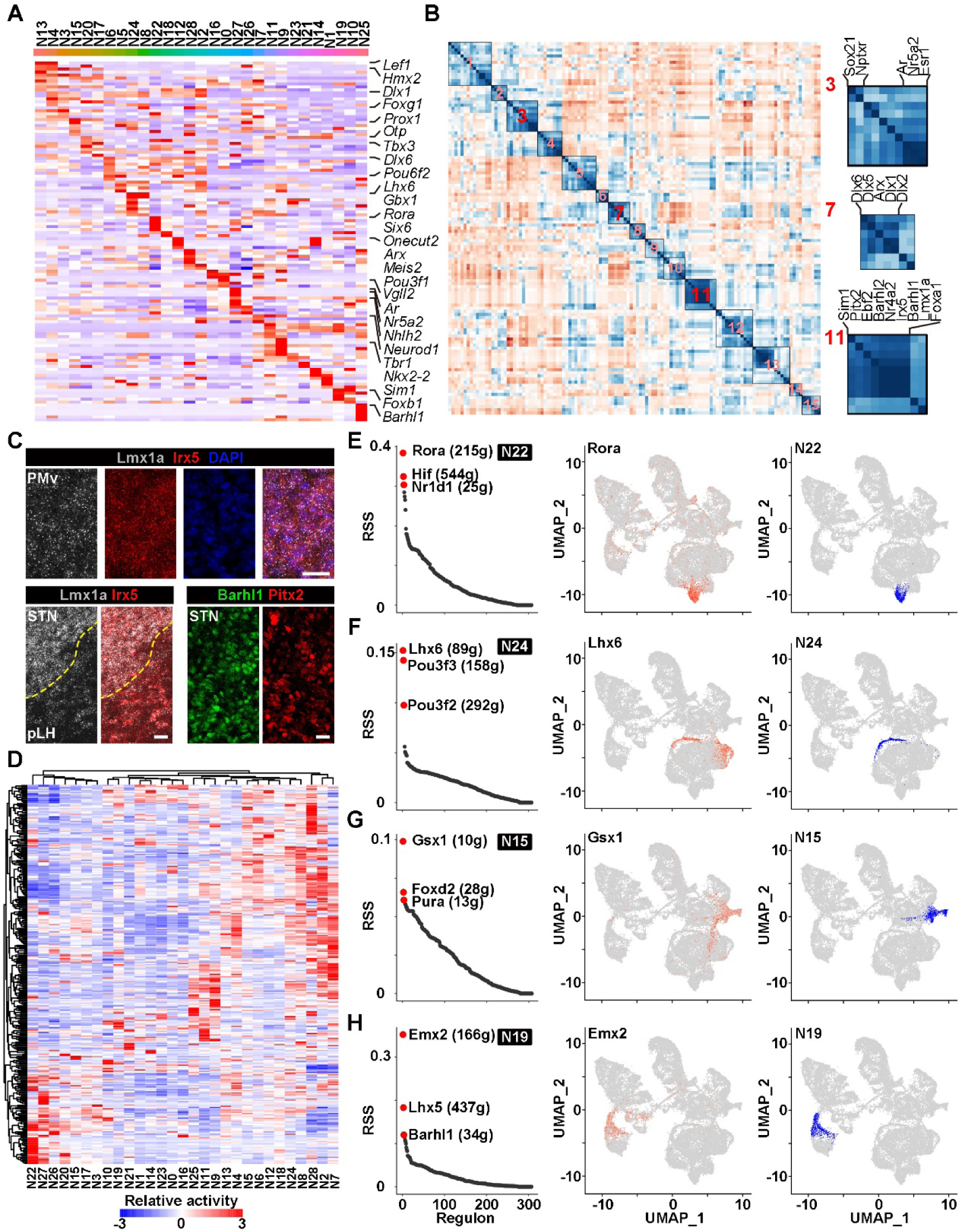
Potential Molecular Determinants of Neuronal Subtype Identity. (A) Shown is the heatmap of 191 subtype-specific TFs expressed in hypothalamic neurons. (B) Matrix showing the 15 TF modules with correlated expression pattern. Modules 3, 7 and 11 with potential regulatory interactions are highlighted. (C) Sample confocal images showing the colocalization of Lmx1a and Irx5 mRNA transcripts and the costaining between Barhl1 and Pitx2 proteins. pLH, posterior lateral hypothalamus; STN, subthalamic nucleus. Scale bars, 20 µm. (D) Heatmap showing the activity of 313 essential regulons for each neuronal subtype. (E-H) Shown are the ranked regulons for N22 (E), N24 (F), N15 (G) and N19 (H) neuronal subtypes, UMAP projection of top regulon expression and UMAP visualization of respective neuronal subtypes (from left to right). The top 3 regulons for each neuronal subtype are labeled and the number of predicted target genes are shown in the brackets. Rora, Lhx6, Gsx1 and Emx2 genes are identified as the top regulons for N22, N24, N15 and N19 subtypes, respectively.

### Fate Diversification into Peptidergic Neurons During Maturation Process

Hypothalamic neurons were traditionally classified based on fast neurotransmitters (e.g. glutamate and GABA), monoamines (e.g. dopamine) and neuropeptides, and featured by a complex neuronal subtype diversity (Hokfelt et al., 2000; Puelles et al., 2012). Here we found that while neuropeptides Gal, Npy, Agrp, Ghrh, Penk and Pnoc were specifically enriched in GABAergic neurons, glutamate could be co-released with Pomc, Cbln1, Tac2, Cck, Reln, Trh, Crh and Adcyap1 (Figure 4E). To provide a systematic view of peptide expression, we examined a total of 100 neuropeptides and monoamines, and identified 34 of them showing specific expression pattern in diverse neuronal subtypes (Figure 6A; Table S9). Further correlation analysis revealed 8 co-expression modules at the cluster level (Figure 6B), implicating the co-release of multiple peptides and monoamines from a single neuronal subtype or the common lineage origin of multiple peptidergic neurons. To test this, we focused on three prominent peptide modules (M3, M4 and M7), evaluated their expression pattern and performed pseudotemporal analysis of representative neuronal subtypes covering both nascent and mature neurons. Our data showed three patterns of peptidergic neuron development: 1) N15 subtype eventually developed into neurons co-releasing Gal, Ghrh and dopamine in arcuate nucleus (ARC), suggesting its fate restriction to a single subtype of peptidergic neurons (Figures 6C and 6D); 2) N20 subtype differentiated into either Agrp/Npy neurons or Sst neurons in ARC, suggesting its fate bifurcation (Figure 6E); and 3) nascent neurons in N21 subtype diversified into Avp, Crh and Oxt neurons in paraventricular nucleus (PVN) during their maturation process, suggesting its multifurcation in cell fate decisions (Figure 6F). Further analysis of N3, N13, N14, N16, N25 and N26 neuronal subtypes confirmed the potential fate diversification of relatively nascent neurons (Figures S12). Together, our results support that hypothalamic peptidergic neuronal subtypes transit through a postmitotic transcriptionally immature state to develop a more complex neuronal diversity.

**Figure 6.**
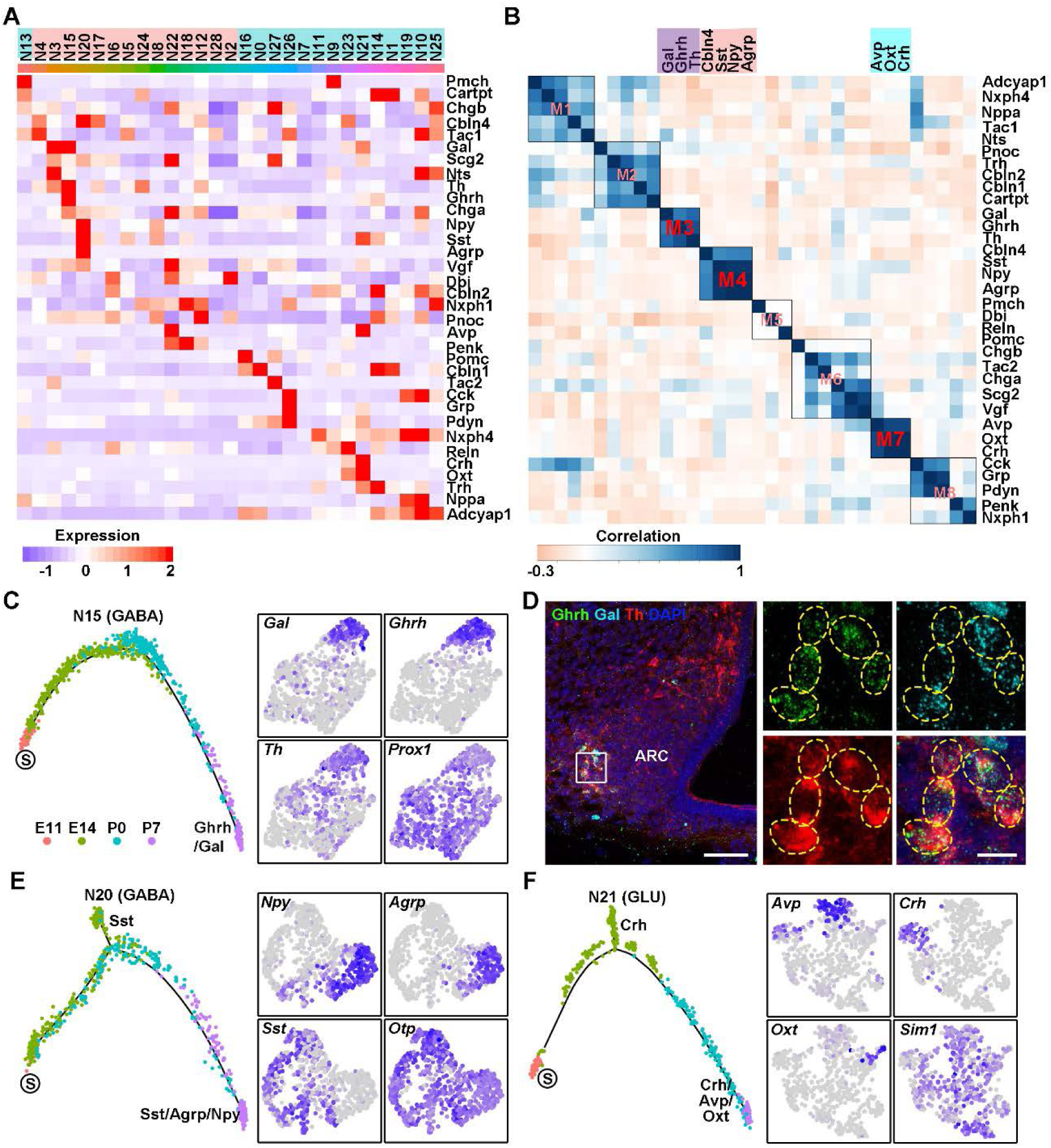
Developmental Diversification of Hypothalamic Peptidergic Neurons (A) Heatmap showing 34 subtype-specific neuropeptides/monoamine expressed in hypothalamic neurons. (B) Matrix showing the 8 neuropeptide/monoamine modules with correlated expression pattern. M3 (Gal, Ghrh and Th), M4 (Cbln4, Sst, Npy and Agrp) and M9 (Avp, Oxt and Crh) modules display a significant correlation at the cluster level. (C) Pseudotemporal trajectory of N15 neuronal subtype specifically expressing M3 neuropeptide modules (left) and the UMAP projection of respective neuropeptide or TF expression (right). S represents starter cells for lineage progression. (D) Sample confocal images showing the co-labeling of Ghrh, Gal and Th in a subpopulation of arcuate neurons within P0 hypothalamus. smFISH was performed to detect *Ghrh* and *Gal* mRNA, followed by immunostaining of TH proteins. Boxed area in the left image is magnified and dashed circles indicate the co-stained hypothalamic neurons. Scale bars, 100 µm (left) and 10 µm (right). (E-F) Pseudotime trajectory of N20 and N21 neuronal subtypes respectively expressing M4 and M9 neuropeptide modules (left) and the UMAP projection of respective neuropeptide expression (right). Otp and Sim1 were selected as the pan-marker for N20 and N21 neuronal subtypes.

## DISCUSSION

Understanding the principles governing the generation of neuronal diversity in complex brain structure and the developmental specification programs that give rise to the diversity is fundamentally important to dissect how neurons are assembled to form functional neural networks. Here, our single-cell transcriptomic analyses of cells derived from Rax^+^ neuroepithelium and spanning multiple developmental stages of hypothalamic neurons suggest a cascade diversifying model in which RGCs, HIPCs and nascent neurons along the cellular hierarchy contribute to the fate diversification of hypothalamic neurons in a step-by-step amplifying fashion, with each step under the precise control of subtype-specific TFs. Our model supports the common developmental principle that the generation of neuronal diversity relies on temporal and spatial patterning of neural progenitors (Holguera and Desplan, 2018; McConnell, 1991; Telley et al., 2019), but differs from the fate-predetermined model wherein cortical RGCs undergo asymmetric cell divisions to produce fate-specified IPCs with a specific spatiotemporal coding and the stochastic model in which retinal progenitor cells (RPCs) are subject to stochastic factors controlling neuronal fate specification (Figure 7) (He et al., 2012; Kohwi and Doe, 2013). Given the extreme neuronal diversity in hypothalamus, it is reasonable for hypothalamic progenitors to adopt a cascade diversifying strategy to yield extraordinary heterogeneity. A very recent study provided an invaluable dataset of developing hypothalamus and a set of regulons shaping the neuronal diversity (Romanov et al., 2020). However, this work did not capture the early RGCs and IPCs for studying their lineage progression due to the lack of collecting cells prior to the peak of hypothalamic neurogenesis at E12.5 (Ishii and Bouret, 2012). We not only uncovered the production of two IPC subpopulations by hypothalamic RGCs and the generic fate-bifurcation of Ascl1^+^ HIPCs, but also analyzed the fate diversification of immature neurons during development, which collectively support the cascade diversifying model.

**Figure 7.**
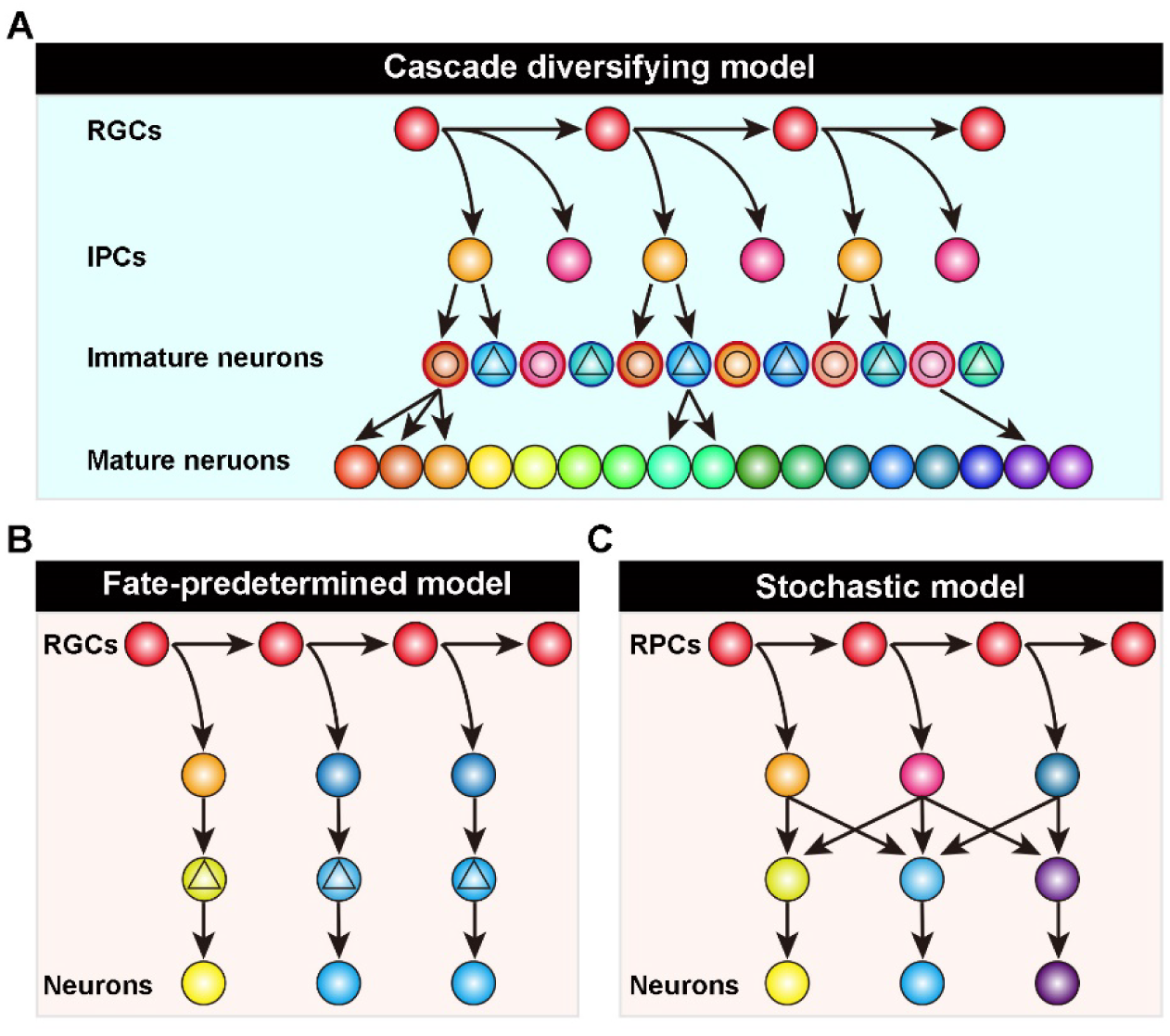
Models for Developmental Origin of Neuronal Diversity. (A) In our “cascade diversifying” model, RGCs, IPCs and nascent neurons along the cellular hierarchy collectively contribute to the generation of neuronal diversity in a stepwise amplifying manner: RGCs generate Ascl1+ and Neurog2+ IPCs; Ascl1+ IPCs produce both GABAergic (spheres with circles) and glutamatergic (spheres with triangles) neurons; and nascent neurons resolve and diversify their fate during maturation. (B) In the “fate-predetermined” model, cortical RGCs sequentially produce fate-restricted Tbr2+ IPCs which subsequently differentiate into deep-(yellow spheres) or upper-layer (blue spheres) excitatory neurons. (C) In the “stochastic” model, retinal progenitor cells (RPCs) stochastically adopt diverse cell fates during differentiation.

As the cells residing at the top of neural lineage hierarchy, multipotential RGCs lining the third ventricle exploit a conserved strategy to sequentially produce neuronal and glial lineages. In contrast to the striking regional specification of RGCs in telencephalon wherein dorsal and ventral subdivisions contain different subpopulations of neural progenitors with diverse potentials in neuronal fate decision (Guillemot, 2005; Parras et al., 2002), hypothalamic RGCs exhibit the potentials to produce Ascl1^+^ and Neurog2^+^ IPCs that are respectively fate-determined to differentiate into GABAergic and glutamatergic neurons in the neocortex (Aydin et al., 2019; Fode et al., 2000; Lo et al., 2002). Besides, we showed that RGCs in embryonic hypothalamus exit cell cycle and switch into pri-TC state at early developmental stage, and thus proposed a “state-switching” hypothesis to interpret the embryonic origin of TCs, previously regarded as the third population of postnatal neural progenitors (Robins et al., 2013). Recent studies have provided “set-aside” and “continuous” models to depict the embryonic origin of adult neural stem cells in lateral ventricles and hippocampus, respectively (Berg et al., 2019; Fuentealba et al., 2015). While hippocampal RGCs continuously produce progeny until they transition into a quiescent state postnatally in the “continuous” model, both “set-aside” and our “state-switching” models suggest that RGCs quit dividing at embryonic stage and remain largely quiescent until postnatally. Nevertheless, hypothalamic RGCs switch into quiescent state much earlier than cortical RGCs to produce postnatal neural progenitors, and we are unable to exclude the possibility that RGCs re-enter the cell cycle after quit dividing but continue with slow cycling.

There coexist Ascl1^+^ and Neurog2^+^ IPCs in the developing hypothalamus, constituting the second-tier cells of neural lineage hierarchy and displaying different molecular profiles, lineage potentials and spatial distribution. It is notable that Ascl1^+^ HIPCs are capable of differentiating into both GABAergic and glutamatergic neurons during their lineage progression, in contrast with the restriction of Ascl1^+^ IPCs to GABAergic fate in telencephalon (Parras et al., 2002). Neurons derived from Ascl1^+^ HIPCs traverse the entire hypothalamus and populate multiple nuclei, including ventromedial nucleus (VMH) and ventral premammillary nucleus (PMv) that are predominantly comprised of glutamatergic neurons. Recently, it has been reported that Ascl1 acts via Neurog3 to specify VMH neuronal fate and loss of Ascl1 causes a reduction in the number of VMH neurons (Aslanpour et al., 2020), supporting the bipotency of Ascl1^+^ HIPCs in neuronal specification.

We further identified nascent postmitotic neurons as the third-tier cells contributing to the generation of neuronal diversity in hypothalamus. As compared with the molecular taxonomy of adult hypothalamic neurons (Campbell et al., 2017; Chen et al., 2017), our single-cell dataset covering the developing hypothalamus contains the immature neurons for which their fates are flexible to be restricted, bifurcated or multifurcated in neuronal subtype specification. Given that there is a rapid induction of neuropeptides in hypothalamus during the early postnatal window (between P2 and P10) (Romanov et al., 2020), we combined pseudotemporal analysis with peptide expression to predict the fate specification of immature neurons among different neuronal sublineages and found that neurons at nascent state are capable of resolving into multiple peptidergic neuronal subtypes during development. Consistent with our hypothesis that nascent neurons could be fate diversified during the process of maturation, a recent work studying peripheral nervous system demonstrated that the divergent somatosensory neurons transition from a transcriptionally unspecialized state to transcriptionally diverse subtypes (Sharma et al., 2020). Moreover, we also detected a plethora of regulons potentially specifying the fate of diverse hypothalamic neuronal subtypes, which are partially overlapping with the gene regulatory networks identified in a very recent study (Romanov et al., 2020). The identified regulons such as Rora, Lhx6, Gsx1, Six6, Otp, Hmx3, Pou3f2 and Lhx2 have previously been found to regulate hypothalamic neuronal fate commitment and physiological functions (e.g. sleeping, metabolic homeostasis, circadian rhythm and endocrine release) (Alvarez-Bolado, 2019; Xie and Dorsky, 2017).

Taken together, we revealed a stepwise amplifying strategy used by neural progenitors to yield an extreme neuronal heterogeneity in evolutionarily conserved brain structure. The disclosure of spatiotemporal patterning of hypothalamic neurogenesis at single-cell resolution provides key insights into the intrinsic nature of neural progenitors and their progeny to develop highly diverse subpopulations of neurons for networking and functioning, although the role of environmental cues in shaping neuronal diversity requires further exploration. Our study will streamline the future studies to generate individual hypothalamic neuronal subtypes *in vitro*, decipher the functional microcircuits among hypothalamic neurons during development and investigate the pathophysiological mechanisms underlying hypothalamic disease.

## METHODS AND MATERIALS

### Animals and Housing

*Rax-CreER^T2^* (Stock No. 025521), *Ascl1-CreER^T2^* (Stock No. 012882), *Ai14* (Stock No. 007914) mouse strains were obtained from the Jackson Laboratory. Animals were maintained on a 12h light-dark cycle with libitum access to food and water. All animal procedures used in this study were performed according to protocols approved by the Institutional Animal Care and Use Committee at Institute of Genetics and Developmental Biology, Chinese Academy of Sciences.

### Tamoxifen Induction

To trace the hypothalamic cell lineages, we bred *Rax-CreER^T2^* and *Ascl1-CreER^T2^* mice with Ai14 reporter mice, checked the vaginal plug to determine the time of pregnancy and intraperitoneally injected a single dose of tamoxifen (132 mg/kg body weight) into pregnant females at E9 to label Rax^+^ hypothalamic neuroepithelium and at E12 to label Ascl1^+^ IPCs in the embryonic brains. Tdtomato-labeled embryos at E11 and E14 were obtained by cesarean section and perfused intracardially with cold 4% paraformaldehyde (PFA) in phosphate-buffered saline (PBS). To collect postnatal brains, we recovered live embryos at E18-E19 by cesarean section, cared the pups with foster female animals and collected the postnatal mice for further analyses.

### Edu Pulse-Chase Assay

To investigate the birthdate of postnatal tanycytes, we performed pulse-chase assay by intraperitoneally apply a single dose of EdU (200 mg/kg body weight) into pregnant female animals at E11 or E14 to label dividing cells. The pups were anesthetized on ice, intracardially perfused with 4% PFA and dissected to collect the brain tissues at P7 for analyzing the label-retaining cells.

### Single-Cell Isolation and Fluorescence Activated Cell Sorting (FACS)

We isolated single cells from *Rax-CreER^T2^::Ai14* mice as previously described with minor modifications (Moffitt et al., 2018). Briefly, the brain tissues were harvested at E11, E14, P0 and P7, followed by the microdissection of hypothalamic tissues under fluorescence stereomicroscope (Lecia M205, Leica Microsystems, Germany). The microdissected tissues were transferred to a 24-well cell culture plate and treated with papain digestion buffer comprising 10 U/mL Papain (Worthington, LK003178), 200 U/mL DNaseI (Worthington, LK003172), 0.5 U/mL chondroitinase ABC (Sigma, C3667), 0.07% hyaluronidase (Rhawn, R006687), 1× Glutamax (Life Technologies, 35050061), 0.05 mM (2R)-amino-5-phosphonovaleric acid (APV; Thermo Fisher Scientific, 010510), 0.01 mM Y27632 dihydrochloride (Sigma, T9531), and 0.2×B27 supplement (Thermo Fisher Biosciences, 17504044) in Hibernate-E media (Life Technologies, A1247601). After digesting 15-30 min (embryonic brains) or 1-1.5 hrs (postnatal brains) at 37°C, the papain digestion buffer was changed into Hibernate-E buffer containing 1×Glutamax, 0.05 mM APV, 0.2× B27, 0.01 mM Y27632 dihydrochloride and 1% fetal bovine serum (FBS). Then tissues were gently triturated through Pasteur pipettes with finely-polished tips of 600-, 300- and 200-μm diameters and washed once with Hibernate-E buffer to generate single-cell suspension. Subsequently, the dissociated hypothalamic cells were stained with DAPI (0.2 µg/mL) to identify dead cells and subjected to fluorescence-activated cell sorting (FACS; BD FACSAria II flow cytometer, BD Bioscience, U.S.A.) for isolating tdTomato-positive live cells.

### Single-Cell RNA Library Preparation and Sequencing

Single-cell RNA-seq libraries were constructed according to the instructions provided by 10×Genomics accompanying single cell 3’ Library and Gel Bead Kit V3 (10×Genomics, 1000075). Cell suspensions (300-600 living cells per microliter determined by Count Star) were loaded on a Chromium Single Cell Controller (10×Genomics) to generate single-cell gel beads in the emulsion (GEM). Captured cells were lysed to release mRNA which were subsequently barcoded through reverse transcribing individual GEMs (Zheng et al., 2017). Using a S1000TM Touch Thermal Cycler (Bio-Rad, U.S.A.) to reverse transcribe, the GEMs were programed at 53°C for 45 min, followed by 85°C for 5 min and hold at 4°C. The cDNA library was then generated, amplified and assessed for quality control using the Agilent 4200. The single-cell RNA sequencing was further performed on the Illumina Novaseq6000 sequencer.

### Tissue Section Preparation, Edu Staining and Immunohistochemistry

The animals were anesthetized by hypothermia and then transcardially perfused with saline followed by 4% PFA in PBS. Mouse brains were immediately dissected, post-fixed for 4-6 hrs in 4% PFA at 4°C, and subsequently cryo-protected in 20% sucrose in PBS for 12 hrs followed by 30% sucrose for 24 hrs. Tissue blocks were prepared by embedding in Tissue-Tek O.C.T. Compound (Sakura 4583). The brain sections (20-40 μm in thickness) were prepared using a cryostat microtome (Leica, CM3050S), dried for 30 min at room temperature in the dark and stored in −20°C freezer. For immunostaining, the tissue sections were washed with 1×TBS (pH=7.4, containing 3 mM KCl, 25 mM Trizma base and 137 mM NaCl) and pre-blocked with 1×TBS++ (TBS containing 5% donkey serum and 0.3% Triton X-100) for 1 hr at room temperature, followed by incubation with primary antibodies diluted in TBS++ overnight at 4°C. The primary antibodies used in this study included goat anti-Sox2 (R&D; AF2018; 1:250), mouse anti-GFAP (Merk-millipore; MAB360; 1:500), mouse anti-BLBP (Abcam; ab131137; 1:250), rabbit anti-Vimentin (Abcam; ab92547; 1:250), rabbit anti-RFP (Rockland; 600-401-379; 1:1000), mouse anti-Ascl1 (BD; 556604; 1:250), mouse anti-Neurog2 (R&D; MAB3314; 1:500), rabbit anti-TH (Novusbio; NB300-109; 1: 500), sheep anti-Pitx2 (R&D; AF7388; 1:500), rabbit anti-Barhl1(Novusbio; NBP1-86513; 1:500). After primary antibody incubation, the brain sections were washed for 3 times with 1×TBS and incubated with following secondary antibodies for 2 hrs at room temperature: anti-mouse Cy2, anti-rabbit Cy3 and anti-rabbit Cy5 (Donkey; Jackson ImmunoResearch; 1:500).

For EdU labeling, the tissue sections were permeabilized with 0.5% Triton-X100 for 30 min and detected with detection solution containing 5 µM Sulfo-Cy3 azide (Lumiprobe, #C1330), 0.1 M Tris-HCl (pH=7.5), 4 mM copper sulfate and 100 mM sodium ascorbate for 30 min. After staining, sections were coverslipped with mounting medium, air-dried overnight and maintained at 4°C in the dark for further imaging. The brain sections were imaged using Leica SP8 equipped with four lasers (405, 488, 568 and 647 nm).

### Single-molecule Fluorescent in Situ Hybridization (smFISH)

To prepare tissue sections for smFISH, mouse brains were dissected, immersed in 4% PFA for 4-6 hrs and then dehydrated with 20-30% DEPC-treated sucrose for 24 hrs. Subsequently, the tissues were rapidly frozen using dry ice, embedded in O.C.T. compound, cryosectioned at a thickness of 30 µm and mounted onto SuperFrost Plus microscope slides. The probes targeting against Slc17a6 (Advanced Cell Diagnostics, #319171) and Slc32a1 (Advanced Cell Diagnostics, #319191) were designed and validated by Advanced Cell Diagnostics. RNAscope v2 Assay (Advanced Cell Diagnostics, #320511) was used for all smFISH experiments according to the manufacturer’s protocol (Wang et al., 2012). Briefly, the brain sections were dried at 55°C for 2 hrs, rinsed with 1×PBS, treated with 3% hydrogen peroxide in methanol and subjected to antigen retrieval. Subsequently, the tissue sections were dehydrated with 100% ethanol and incubated with mRNA probes for 2 hrs at 40°C. The specific signals were then amplified with multiplexed amplification buffer and detected with TSA Plus Fluorescence kits (Perkin Elmer, #NEL753001KT).

To detect the mRNA expression level of Gal, Ghrh, Irx5 and Lmx1a, we used hybridization chain reaction (HCR) approach (Choi et al., 2018). A software (available at https://github.com/GradinaruLab/HCRprobe) was used to design HCR probes for targeting coding sequence (CDS) and 3’UTR region of interest gene genes (Patriarchi et al., 2018). The sequences of all HCR probes are included in Table S10 and all of them were synthesized by Sangon Biothch, China. The brain sections were permeabilized in 70% ethanol for 16 hrs at 4°C, followed by 0.5% Triton X-100 in 1×PBS at 37°C for 1 hr, and treated with 10 µg/mL Protease K was to improve mRNA accessibility. After two washes with 1×PBS at room temperature, sections were pre-hybridized in 30% probe hybridization buffer for 1 hr at 37°C and then incubated in 30% probe hybridization buffer containing HCR probes (10 µM for each) at 37°C for 3 hrs. After mRNA hybridization, the washing and amplification steps were performed as previously described (Choi et al., 2018). Immunofluorescence staining with antibodies against TH was further conducted on HCR-labeled sections.

### Cross-Reference of Allen Developing Mouse Brain Atlas

*In situ* images of mouse brains were resourced from Allen Developing Mouse Brain Atlas (http:// developingmouse.brain-map.org). To map the spatial distribution of HIPC subtypes, we marked the peduncular and terminal regions along longitudinal axis, as well as preoptic, anterior, tuberal and mammillary zones along sagittal axis on the E13.5 reference sections. To identify the nucleus distribution of diverse neuronal subtypes, we referred the prenatal atlas and marked diverse hypothalamic nuclei on the E18.5 and P4 reference sections.

### Pre-Processing of Sequencing Data

Read pre-processing was performed using the 10×Genomics workflow. Briefly, the Cellranger’s pipeline (version 3.0.2) was used for demultiplexing raw sequencing data, barcode assignment and quantification of unique molecular identifiers (UMIs). Using a pre-built reference package, we mapped the reads to the mouse reference genome (GRCm38/mm10, release 93) to generate gene expression matrices. The matrix files arising from different batches of experiments were subsequently analyzed using Seurat v3 software (Stuart et al., 2019).

### Quality Control, Data Filtration and Integration of Datasets

For quality control, we set the threshold and removed cells with more than 5% of reads mapping to mitochondrial genes (considered to be low-quality cells that exhibit extensive mitochondrial contamination), fewer than 2,000 genes (considered to be low-quality cells or empty droplets), more than 8,000 genes or 50,000 UMIs (considered likely to be cell doublets or multiplets) for subsequent analyses. Moreover, we manually excluded the contaminating haemoglobin genes (*Hbb-bt*, *Hbb-bs*, *Hbb-bh2*, *Hbb-bh1*, *Hbb-y*, *Hbs1l*, *Hba-x*, *Hba-a1*, *Hbq1b*, *Hba-a2* and *Hbq1a*) introduced by the lysis of erythrocyte to avoid any bias, leaving 28,868 genes in 43,670 cells for further integrated analysis.

We integrated multiple datasets covering hypothalamic cells across four different time points using the standard integration workflow built in Seurat v3 software according to the tutorial at https://satijalab.org/seurat/v3.1/integration.html. In brief, logarithmic normalization was implemented separately for each dataset, 2000 variable feature genes were selected and the ‘anchors’ identified by ‘FindIntegrationAnchors’ function were used to integrate data.

### Cell-Cycle Analysis

Variations in cell cycle stages, particularly among mitotic cells transitioning between S and G2/M phases, can lead to a substantial mask of authentic biological signals and thereby alter the cell clustering. Given that neural progenitor cells in the developing brain are mitotically active, we regressed out cell cycle variation to mitigate its adverse effect for clustering RGCs and IPCs by rescaling the score of 43 S-phase and 54 G2/M-phase genes as described previously (Tirosh et al., 2016). Briefly, we assigned cell cycle score for each cell using supervised analysis of the phase-specific genes, calculated the difference between G1/S and G2/M phase scores and subtracted the source of heterogeneity from our datasets to remove cell cycle effect.

### Dimensionality Reduction, Clustering and Visualization

With integrated expression values in hand, we detected the highly variable genes, performed principle component analysis (PCA) and selected the top 40 significant principle components (PCs) to perform Uniform Manifold Approximation and Projection (UMAP) analysis. Using UMAP algorithm, we reduced variation to two dimensions by ‘RunUMAP’ function and carried out unsupervised clustering of cells by constructing a shared nearest neighbor (SNN) graph according to K-nearest neighbors (K=20) and then determining the number of clusters with a modularity function optimizer. The clusters were further compared pairwise to identify cell type-specific genes and the cell identities were assigned by cross-referencing their marker genes with known neural subtype markers. A small subpopulation of cells that ambiguously expressed multiple marker genes for different cell types was found to capture low level of UMIs and defined as low-quality cells or multiplets that escaped the first quality-control step and thereby excluded from further analyses. A final dataset of 43261 individual cells was preserved to map the landscape of hypothalamus development and roughly classified into radial glial cells (RGCs), intermediate progenitor cells (IPCs), excitatory glutamatergic neurons, inhibitory GABAergic neurons, astrocytes, oligodendrocyte progenitor cells (OPCs), oligodendrocytes and ependymal cells. For subclustering, RGCs, IPCs and neurons were separately subsetted and reclustered with an optimal resolution based on the number of cells according to the Seurat guidelines. A dendrogram showing the relatedness of cell subclusters was generated for 29 neuronal subtypes based on a distance matrix constructed in PCA space (BuildClusterTree function).

### Differential Gene Expression Analysis and Functional Annotation

To identify genes differentially expressed in cells of diverse types or at different developmental stages, we compared cell groups pairwise and identified differentially expressed genes using ‘FindAllMarkers’ function with Wilcoxon rank-sum test, which returned logarithmic fold-changes of the average gene expression among groups and adjusted *p* values for each tested gene based on Bonferroni correction. The top-ranking differentially expressed gene sets were chosen for further gene ontology (GO) enrichment analysis with clusterProfiler software (Yu et al., 2012). Functional enrichment of target gene sets in GO biological processes or cellular components was determined with hypergeometric test, setting *p* value threshold of 0.01 for statistical significance. Heatmaps of the differentially expressed genes were generated by ggplot2 package built with Seurat software.

### Monocle Pseudotemporal Analysis

We performed pseudotemporal analysis using Monocle software to infer the developmental trajectory of hypothalamic cell lineage. The Monocle 3 pipeline (Cao et al., 2019) was applied to reconstruct the single-cell trajectories of hypothalamic cells covering neural progenitors and postmitotic neurons, given its optimized designing for large datasets and powerful ability to learn very complex, potentially disjoint trajectories. Briefly, we normalized the expression value by logarithmic transformation and assigned UMAP coordinates from the integrated Seurat object to ‘cell_data_set’ (CDS) object produced by Monocle 3 for trajectory inference.

To simply reconstruct individual trajectories for neuronal and glial lineage, we subsetted neurons or glial cells to integrate with their progenitor cells and performed trajectory analysis using Monocle 2 (Qiu et al., 2017; Trapnell et al., 2014). To order the genes along pseudotime, we imported 2000 variable features identified by Seurat, filtered out genes detected within lower than 5% of cells, applied ‘DDRTree’ to reduce dimensions and plotted the minimum spanning tree of cells. We then clustered genes by their pseudotemporal expression pattern and generated the heatmaps to visualize the transcriptional programs that co-vary across pseudotime. The trends of pseudotemproal gene expression for each transcriptional program were analyzed as previously reported (Xiao et al., 2018).

### Gene Expression Signature Scoring

Individual neuronal subtypes were scored for HIPC signature programs to infer the transcriptional connection between HIPC and neuronal subtypes. Briefly, the signature programs of HIPC subtypes were derived from the top-ranking variable genes identified by the ‘FindAllMarkers’ function with an ‘min.pct’ of 0.25 and ‘logfc.threshold’ of 0.25. We then scored individual neuronal subtypes for their enrichment of HIPC gene signatures using the function ‘AddModuleScore’ in Seurat.

### RNA Velocity Analysis

RNA velocity approach can estimate the cell state by distinguishing between unspliced and spliced mRNAs in single-cell dataset, predict the future state of individual cells and infer their developmental trajectory (La Manno et al., 2018). We performed RNA velocity analysis following the guidance of velocyto.R tutorial at http://velocyto.org/velocyto.py/citing/index.html. The output BAM files derived from Cellranger-based read mapping were used as an input to calculate gene-relative velocity using K-nearest neighbor cell pooling (K=25) and a stationary solution (deltaT=1). We visualized RNA velocities as a regular grid using linear scaling on the UMAP representation generated in Seurat.

### CytoTRACE Analysis

We leveraged CytoTRACE (cellular trajectory reconstruction analysis using gene counts and expression) approach, a recently developed computational method for predicting the direction of differentiation from scRNA-seq data across multiple developmental stages (Gulati et al., 2020), to recover the differentiation and maturation state of neurons in our dataset. The multiple single-cell datasets covering neurons and their immediate progenitors were re-integrated using ‘iCytoTRACE’ function and used to quantify the number of expressed genes as a simple, yet robust indicator of developmental potential. We further visualized the maturation state of neurons on the UMAP embeddings generated previously in Seurat.

### TF and Neuropeptide Modular Analysis

We firstly assessed the expression of 4482 TFs and 100 peptides in diverse neuronal subtypes and identified 191 TFs and 34 neuropeptides with subtype-specific expression pattern. To determine the TF and peptide modules that define potential gene regulatory network and neuronal subtype identities at the cluster level, we calculated the Pearson’s correlation coefficients of subtype-specific TFs and neuropeptides/monoamine based on their averaged gene expression level in 29 neuronal subtypes. The correlation matrices were then imported to corrplot R package (https://github.com/taiyun/corrplot) for a graphical display.

### Regulon Identification for 29 Neuronal Subtypes

Regulon scores for individual cells were computed using the SCENIC (single-cell regulatory network inference and clustering) pipeline (Aibar et al., 2017). A log-normalized expression matrix of neuronal cells was used as an input into the pySCENIC workflow (https://pyscenic.readthedocs.io/en/latest/index.html) with default settings to infer regulons (master TFs and their target genes). Firstly, the potential TF targets were inferred based on our gene expression dataset using the ‘grnboost2’ function. Secondly, we identified a list of enriched TF-binding motifs and the corresponding target genes for all gene co-expression modules, and uncovered the regulons based on the database containing motifs with genome-wide rankings. Thirdly, the activity of each regulon across the cells was scored using the area under the recovery curve (AUC) threshold of genes that define this regulon. Lastly, average regulon activity for 313 identified regulons in 29 neuronal subtypes was computed and visualized as heatmap, which showed groups of regulons that tend to be active in each neuronal subtype.

To quantify the cell-type specificity of a regulon, we converted regulon activity scores (RAS) to regulon specificity scores (RSS) based on the Jensen-Shannon Divergence (JSD) as previously described (Cabili et al., 2011; Suo et al., 2018), followed by the ranking of 313 regulons based on their RSS scores for each neuronal subtype.

### Data Availability

The single-cell RNA-seq data from this study have been deposited in the Gene Expression Omnibus (GEO) with accession number GSE151060. All other relevant data that support the findings of this study are available from the corresponding authors upon reasonable request.

### Code Availability

The computational code used in this work is available upon request.

## ACKNOWLEDGMENTS

We thank Yinqing Li, Samuel Wong, Xiaohong Xu and Jie He for comments on the manuscript, and Lan Jiang for intensive discussions. We gratefully acknowledged Zhengang Yang for providing us *Ascl1-CreER^T2^* mice for lineage tracing. Funding: The work was supported by The Ministry of Science and Technology of China (2019YFA0800213 and 2018YFA0801104), National Natural Science Foundation of China (31771131 and 81891002), Strategic Priority Research Program of Chinese Academy of Sciences (XDB32020000), Hundred-Talent Program (Chinese Academy of Sciences) and Beijing Municipal Science & Technology Commission (Z181100001518001).

## AUTHOR CONTRIBUTIONS

Q.W. and Y.Z. conceived of the presented idea, designed the experiments and developed the theory. M.X., Y.Z. and X.G. performed single-cell isolation for RNA sequencing. Y.Z. and H.W. performed the bioinformatic analysis. M.X., S.L., W.M., X.G. and Q.W. performed immunofluorescence, *in situ* hybridization and imaging. L.G. and M.H. contributed to lineage tracing. Q.W. and Y.Z. wrote the manuscript.

## Supplemental Information

### Figures and Figure Legends

**Figure S1.**
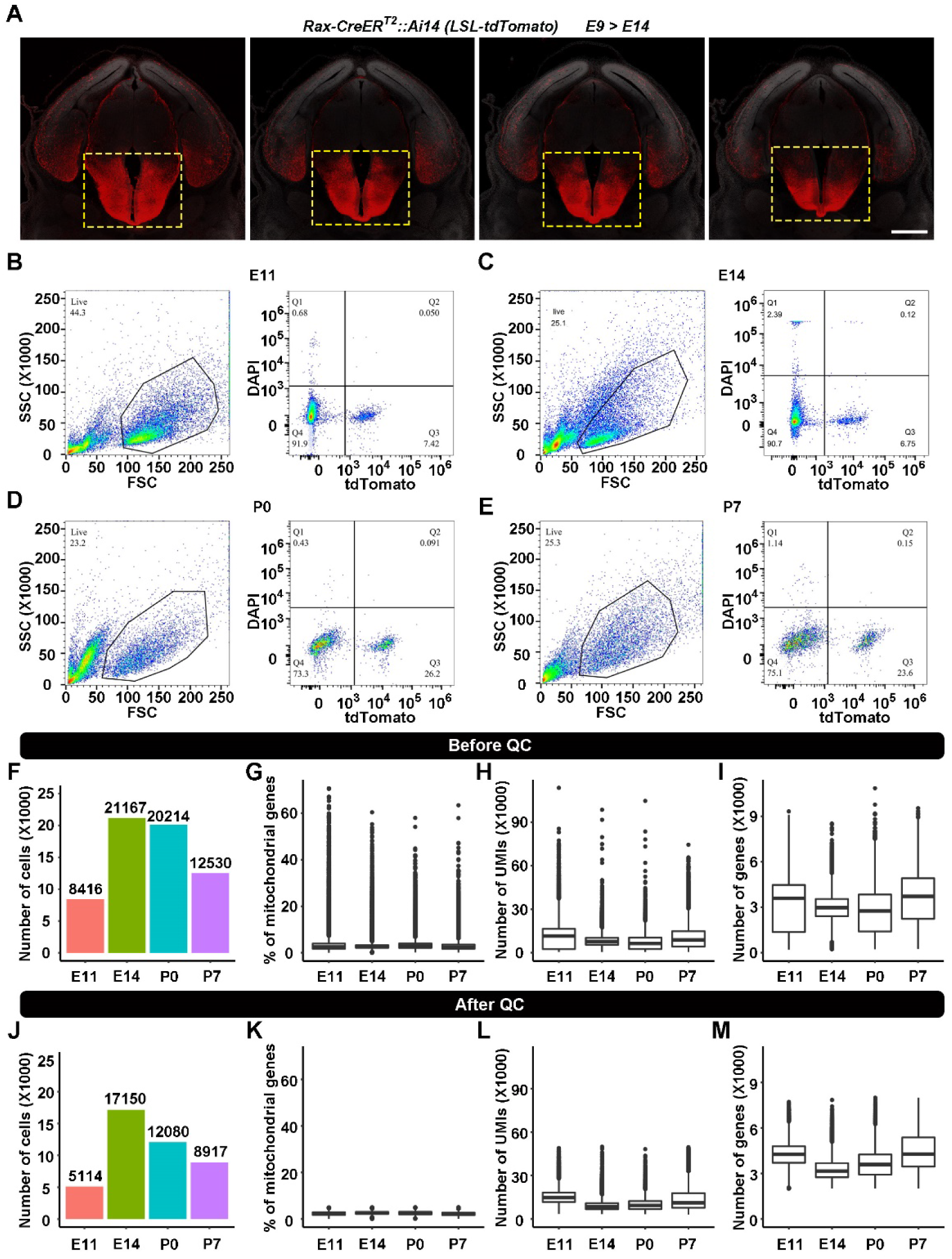
Single-Cell RNA Sequencing and Quality Control Metrics for Developing Hypothalamic Cells. Related to Figure 1. (A) Representative image showing the genetically-mediated fluorescent labeling of hypothalamic cell lineage with *Rax-CreER^T2^::Ai14* mice. The mice were induced with tamoxifen at a single dose of 132 mg/kg body weight at E9 and collected at E14. Scale bar, 500 µm. (B-E) Fluorescence-activated cell sorting (FACS) analysis of tdTomato+ cells from *Rax-CreER^T2^::Ai14* mice induced with tamoxifen at E9 and collected at E11 (B), E14 (C), P0 (D) and P7 (E) for single-cell RNAseq. (F) Bar plot showing the number of sequenced cells collected at different time points. (G-I) Quantification of the percentage of mitochondrial genes, number of UMIs and number of detected genes per cell before quality control. (J) Bar plot showing the number of qualified cells for further bioinformatics analyses. Individual cells with >5% mitochondrial counts (considered to be low-quality or dying cells that exhibit extensive mitochondrial contamination), fewer than 2,000 genes (considered to be low-quality cells or empty droplets), more than 8,000 genes or 50,000 UMIs (considered likely to be cell doublets or multiplets) were eliminated from subsequent analyses. (K-M) Quantification of the percent of mitochondrial genes, number of UMIs and number of detected genes per cell after cell filtering.

**Figure S2.**
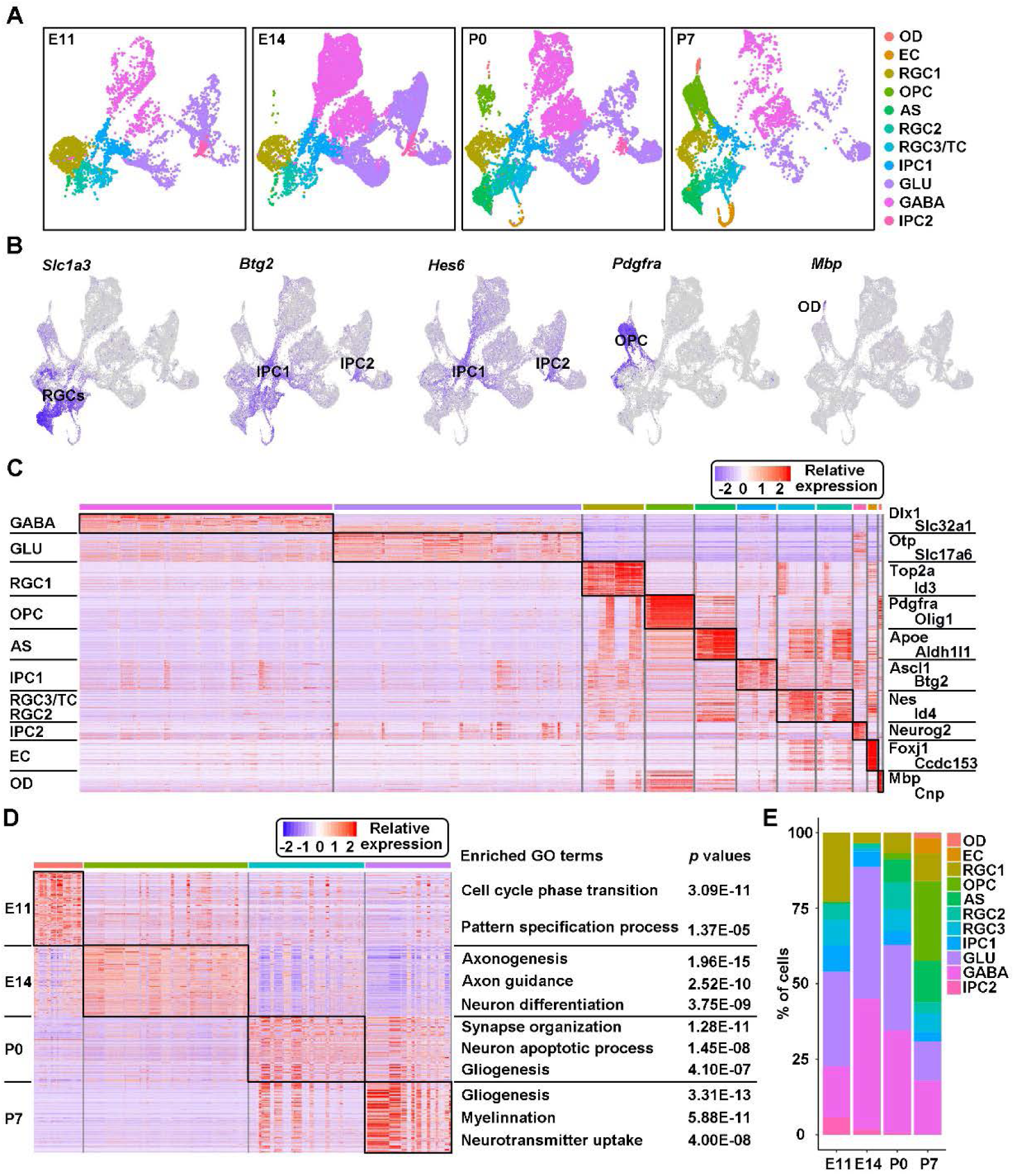
Cell Typing of Developing Hypothalamic Cells. Related to Figure 1. (A) UMAP visualization of diverse hypothalamic cell types at different time points. (B) Imputed gene expression of putative subtype-specific markers overlaid on UMAP representation. (C) Heatmap showing the top 50 differentially expressed genes enriched among diverse hypothalamic cell types. Marker genes defining the identity of each cell type are labeled. (D) Shown is the heatmap of top 100 genes differentially expressed in E11, E14, P0 and P7 hypothalamus. Terms of enriched gene ontology (GO) for each cell type are labeled at the right. (E) Shown is the proportion of diverse hypothalamic cell types at different developmental time points.

**Figure S3.**
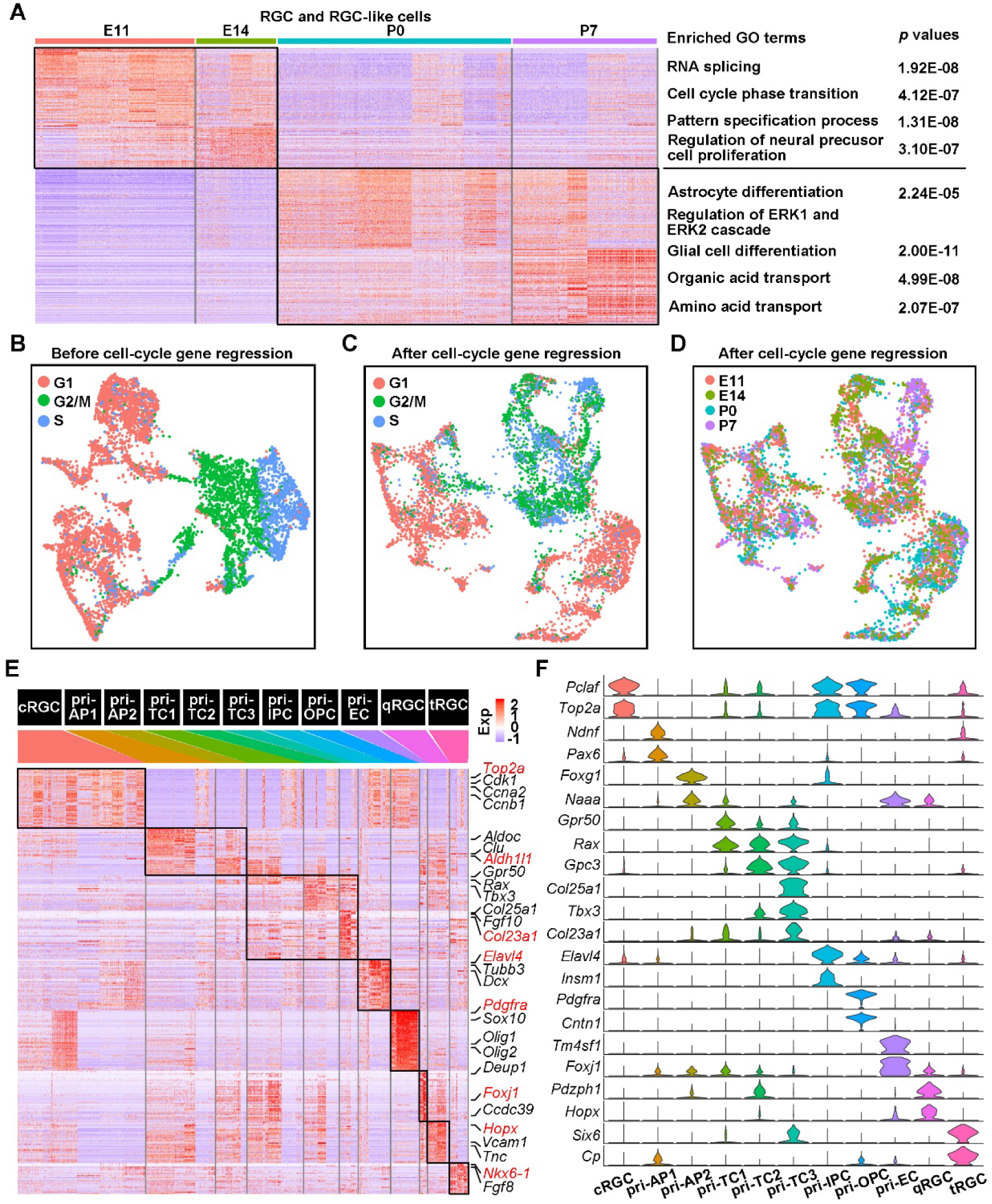
Molecular Landscape of Hypothalamic RGCs. Related to Figure 2. (A) Heatmap showing the top 100 genes differentially expressed in E11, E14, P0 and P7 RGCs or RGC-like cells. The enriched GO terms are shown at the right. (B-C) UMAP visualization of RGCs and RGC-like cells before (B) and after (C) regressing out the effect of cell cycle. The phases of cell cycle are defined by the expression of 43 S-phase and 54 G2/M-phase marker genes. (D) UMAP visualization of 7230 RGCs and RGC-like cells colored by their developmental stages. (E) Heatmap showing the top 50 genes differentially expressed in each RGC subtype. (F) Violin plots showing the expression of RGC subtype-specific genes.

**Figure S4.**
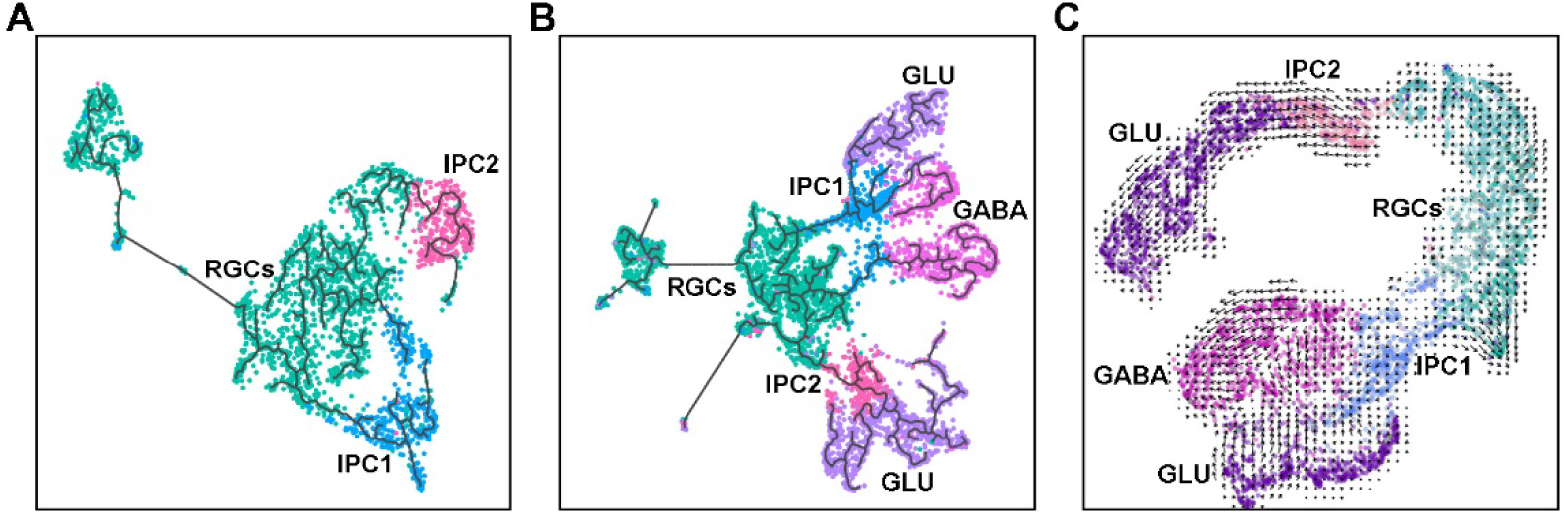
Fate Bifurcation of Hypothalamic RGCs. Related to Figure 2. (A) UMAP visualization of the developmental trajectory of E11 neural progenitors (RGCs and IPCs). The branched trajectory infers the differentiation of RGCs into both IPC1 and IPC2 sublineages. (B) Developmental trajectory of E11 cells within neuronal lineage, including RGCs, IPCs and neurons. (C) RNA velocity of single cells arising from neuronal lineage, overlaid on UMAP representation. The direction of arrows predicts the lineage progression of neural progenitors.

**Figure S5.**
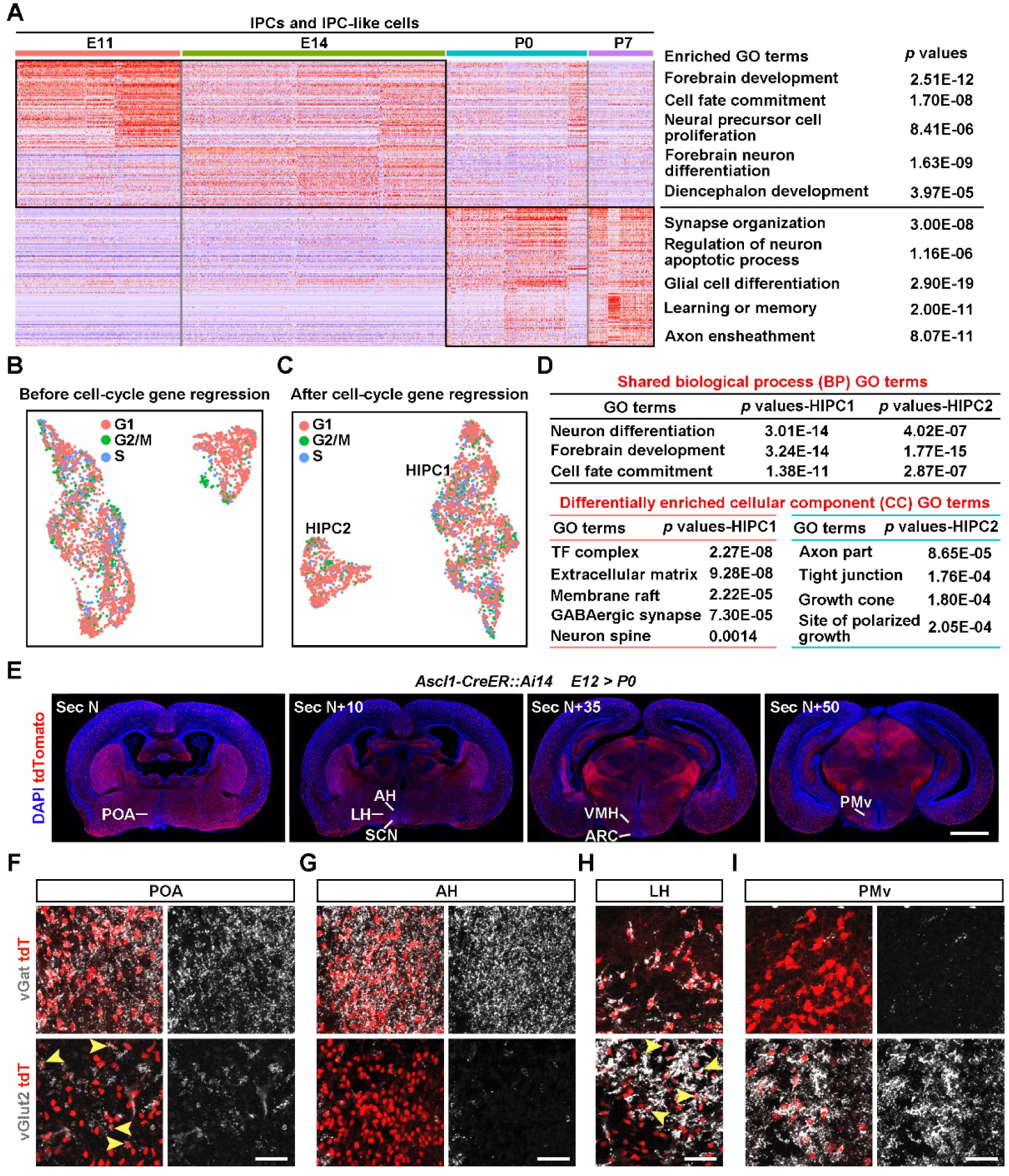
Developmental Landscape and Linage Potential of HIPCs. Related to Figure 3. (A) Heatmap showing the top 100 genes differentially expressed in E11, E14, P0 and P7 hypothalamic IPCs, with enriched GO terms shown at the right. (B-C) UMAP visualization of HIPCs distribution before (B) and after (C) cell-cycle regression. (D) The shared biological process GO terms (top) and different cellular component GO terms (bottom) are shown. (E) Representative images showing the contribution of Ascl1 lineage cells to different brain regions. The *Ascl1-CreER^T2^::Ai14* mice were induced at E12 and sacrificed at P0 for analysis. In the hypothalamus, Ascl1 lineage cells were widely distributed in POA, AH, SCN, LH, VMH, ARC and PMv. POA, preoptic area; AH, anterior hypothalamus; LH, lateral hypothalamic nucleus; SCN, suprachiasmatic nucleus; VMH, ventromedial hypothalamic nucleus; PMv, ventral premammillary nucleus. Scale bar, 1 mm. (F-I) Sample images showing the generation of GABAergic (vGat^+^) and/or glutamatergic (vGlut2^+^) neurons in POA, AH, LH and PMv by Ascl1^+^ HIPCs. Arrowheads indicate the glutamatergic neurons within Ascl1 lineage cells. Scale bars, 50 µm.

**Figure S6.**
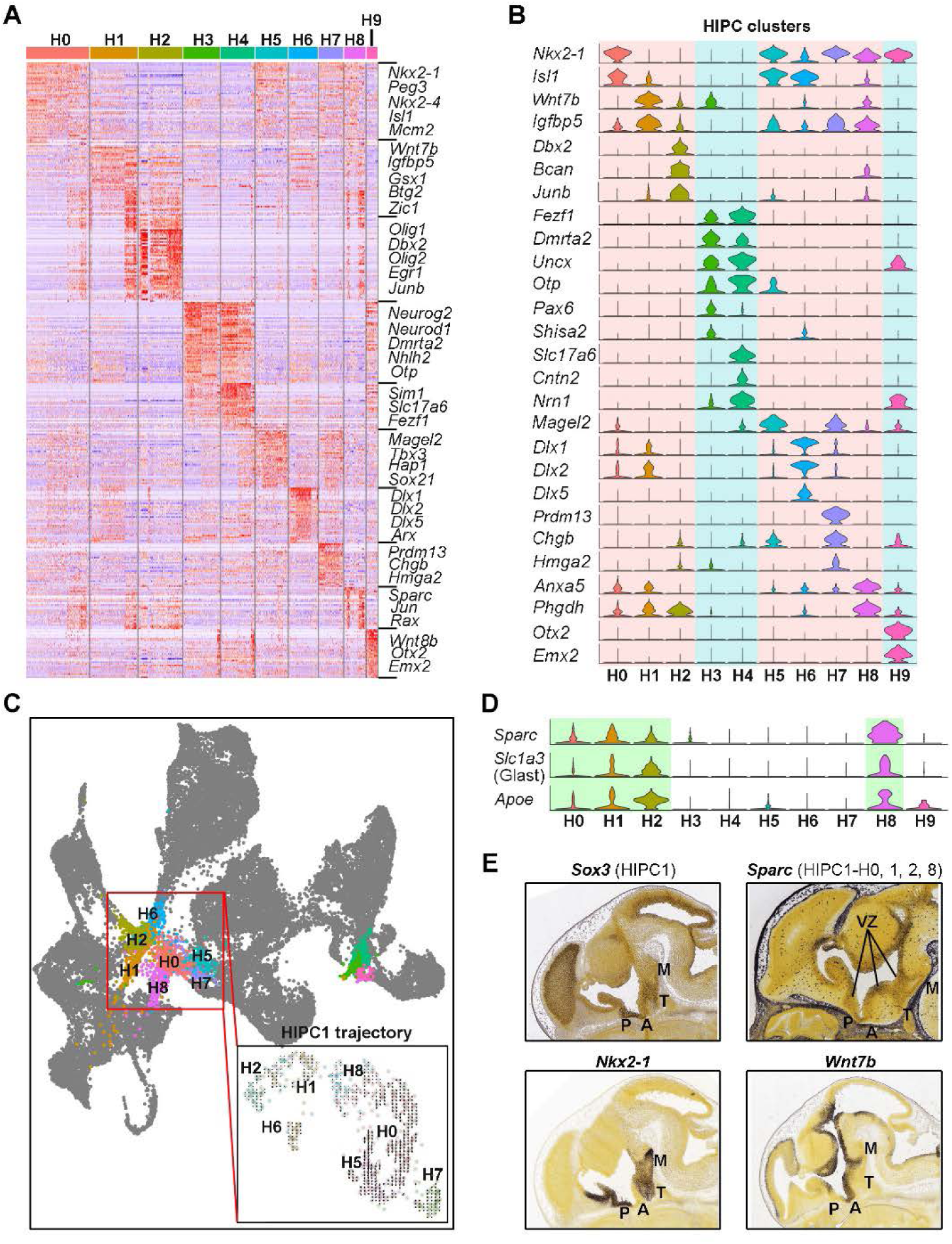
Molecular Profiles and Spatial Distribution of HIPCs. Related to Figure 3. (A) Heatmap showing the top 50 genes differentially expressed in diverse HIPCs subtypes. Specific genes related to each subtype are labeled at the right. (B) Violin plots showing the relatively specific marker genes for each HIPC subtype. (C) UMAP-based trajectory inference of HIPC1 shows a more primitive state of H0-2 and H8 subgroups. Boxed HIPC1 trajectory is inferred by RNA velocity. (D) Violin plots showing the specific expression of stemness genes (Sparc, Slc1a3 and Apoe) in H0-2 and H8 subgroups. (E) Spatial gene expression of Sox3, Sparc, Nkx2-1 and Wnt7b in the E13.5 sagittal brain sections.

**Figure S7.**
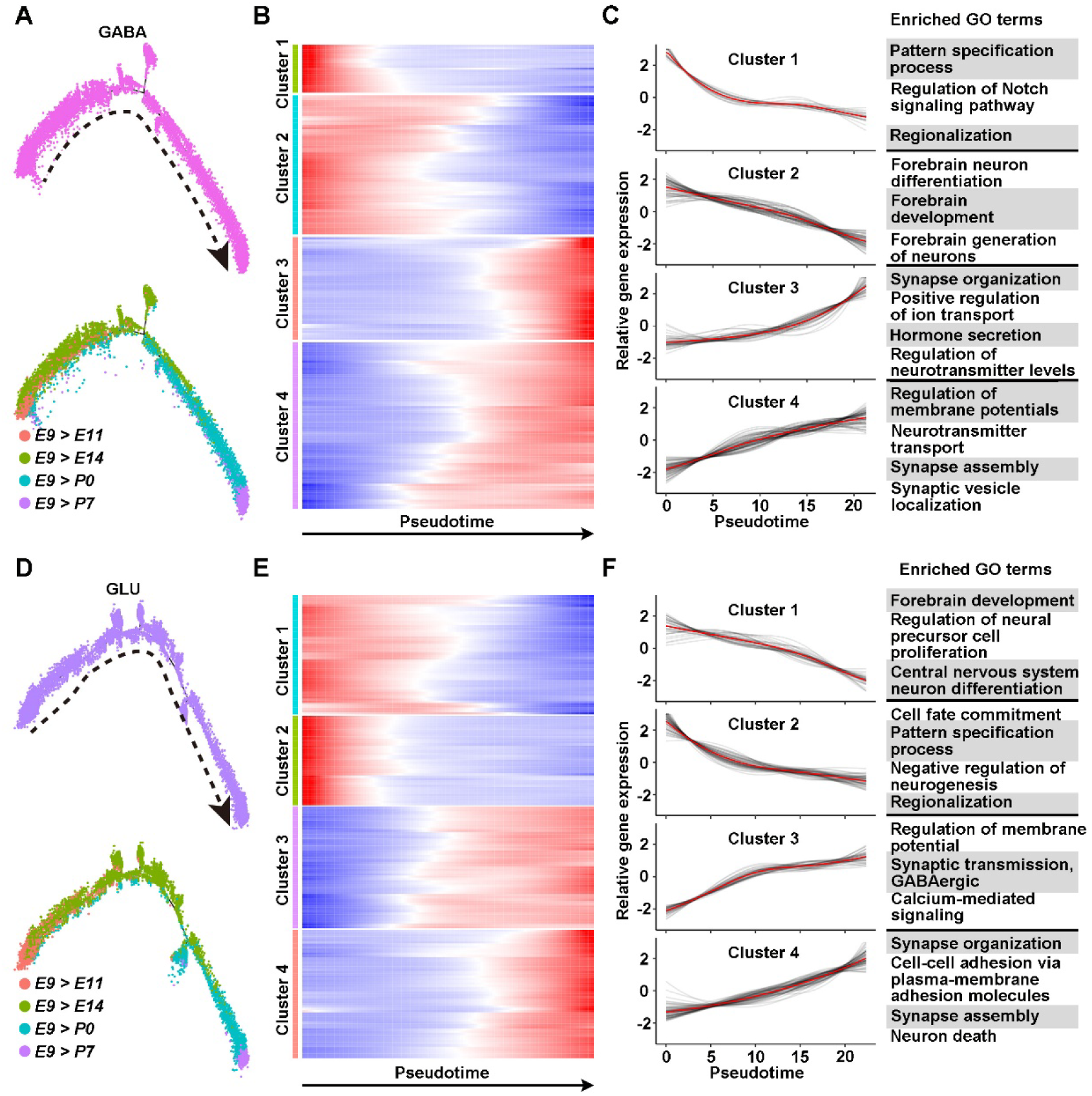
Developmental Progression of Differentiated GABAergic and Glutamatergic Neurons. Related to Figure 4. (A) Pseudotemporal trajectory of GABAergic neurons. The bottom trajectory is colored by the developmental stages of cells. Dashed arrow indicates the direction of developmental progression. (B) Pseudotime-ordered heatmap showing 4 transcriptional programs dictating the developmental progression of GABAergic neurons. The color from blue to red indicates low to high gene expression levels. (C) Shown are gene expression trends for each transcriptional program (left) and their top enriched GO terms (right). The red lines depict the average gene expression level of each transcriptional program, while the gray lines show the temporal expression dynamics for each gene. (D) Pseudotemporal trajectory of glutamatergic neurons. The bottom trajectory is colored by the developmental stages of cells. (E) Pseudotime-ordered heatmap showing 4 transcriptional programs dictating the developmental progression of glutamatergic neurons. (F) Shown are gene expression trends for each transcriptional program (left) and their top enriched GO terms (right).

**Figure S8.**
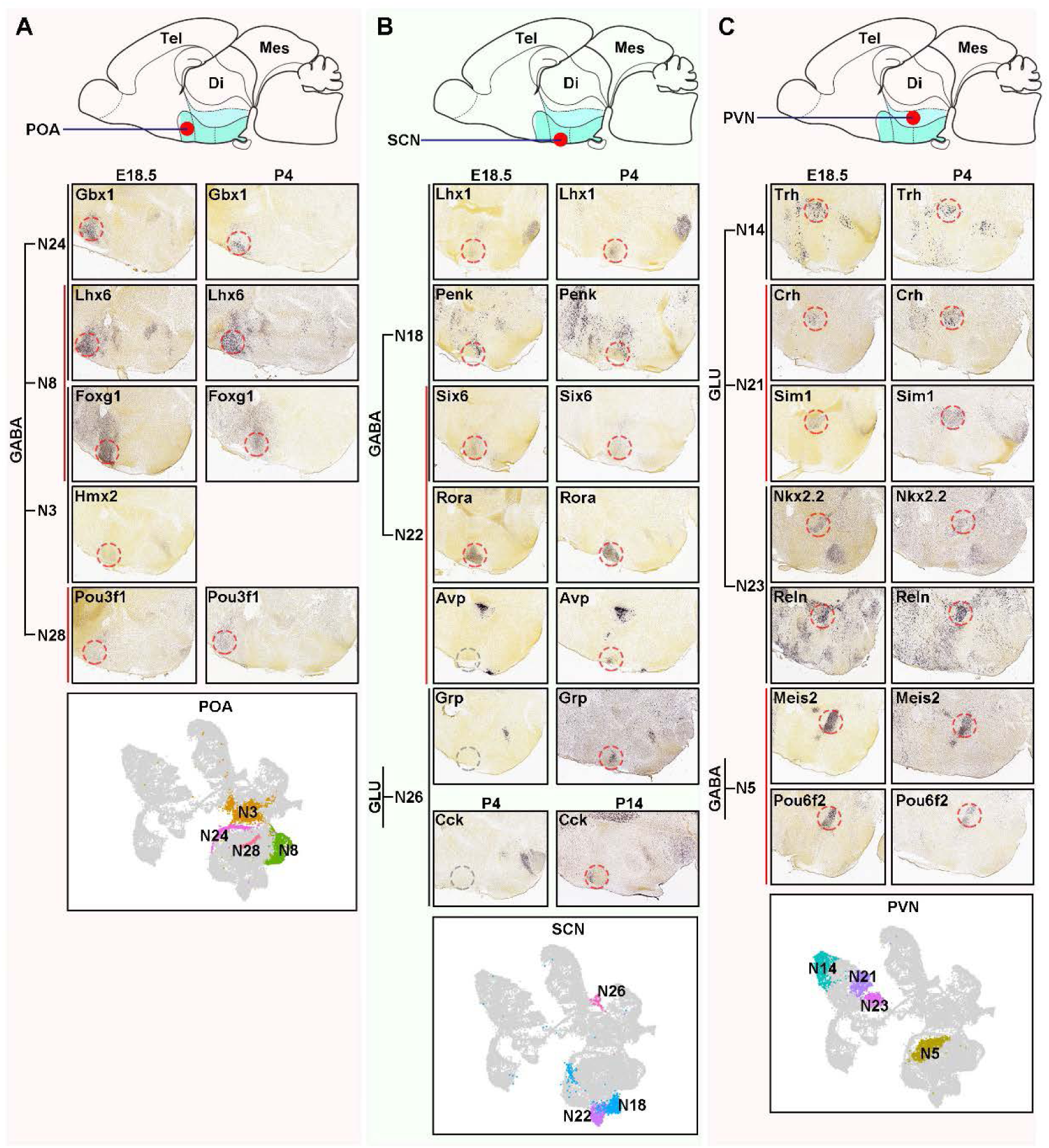
Spatiotemporal Expression of Marker Genes in Several Neuronal Clusters. Related to Figure 4. Shown are *in situ* hybridization layouts of neuronal clusters enriched in POA (A), SCN (B) and PVN (C). Marker genes for each neuronal cluster were chosen for cross-referencing the *in situ* hybridization data in the Allen Developing Mouse Brain Atlas. Neuronal clusters distributed within POA (A), SCN (B) and PVN (C) are also highlighted in boxed UMAP plots (bottom).

**Figure S9.**
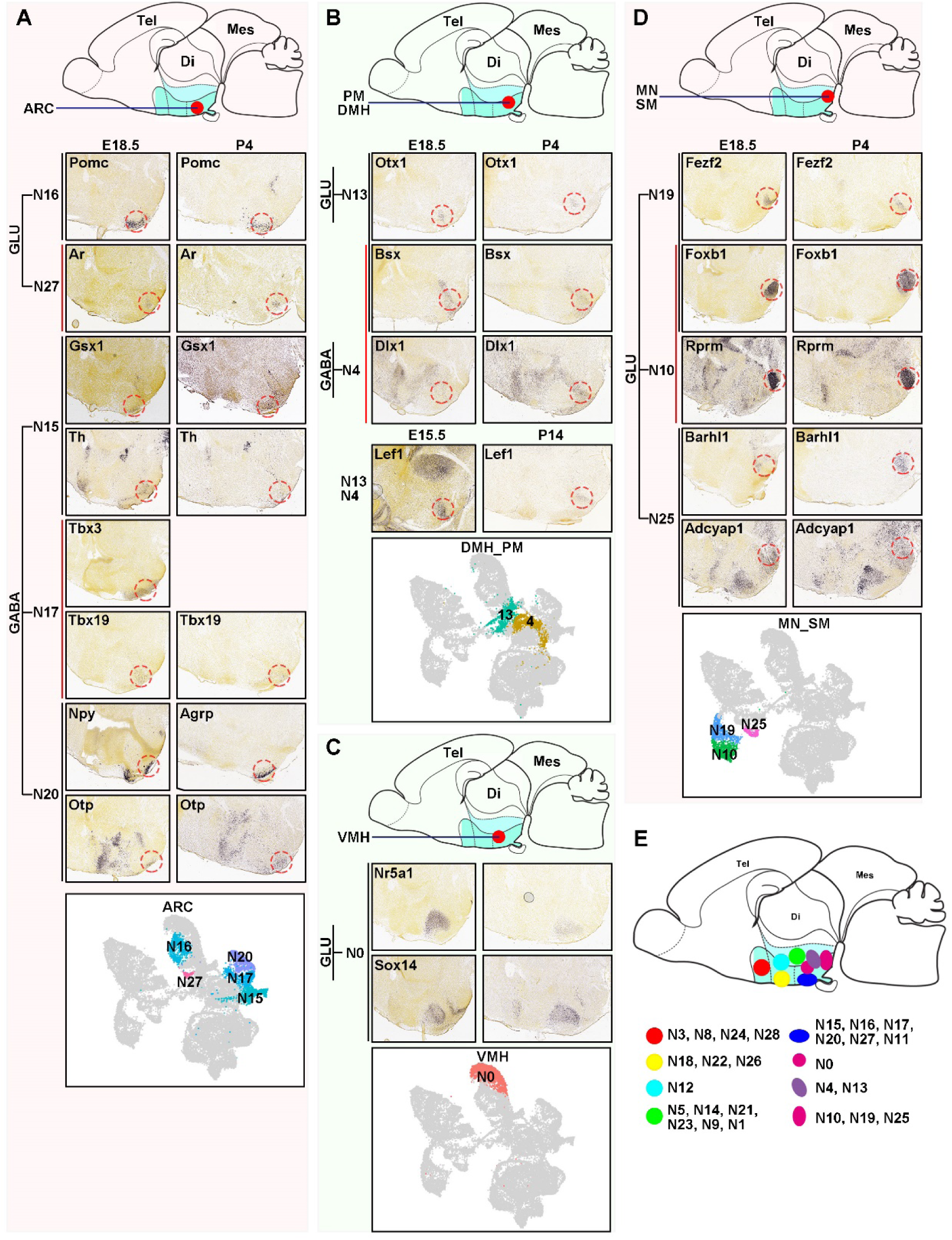
Spatiotemporal Expression of Marker Genes in Several Neuronal Subtypes. Related to Figure 4. (A-D) Shown are *in situ* hybridization layouts of neuronal clusters enriched in ARC (A), DMH/PM (B), VMH (C) and MN/SM (D). Marker genes for each neuronal subtype were chosen for cross-referencing the *in situ* hybridization data in the Allen Developing Mouse Brain Atlas. Neuronal clusters distributed within ARC (A), DMH/PM (B), VMH (C) and MN/SM (D) are also annoated in boxed UMAP plots (bottom). PM, perimamillary nucleus; DMH, dorsomedial hypothalamic nucleus; MN, mammillary nucleus; SM, supramammillary nucleus. (E) Schematic diagram showing the spatial orientation of distinguishable neuronal subtypes.

**Figure S10.**
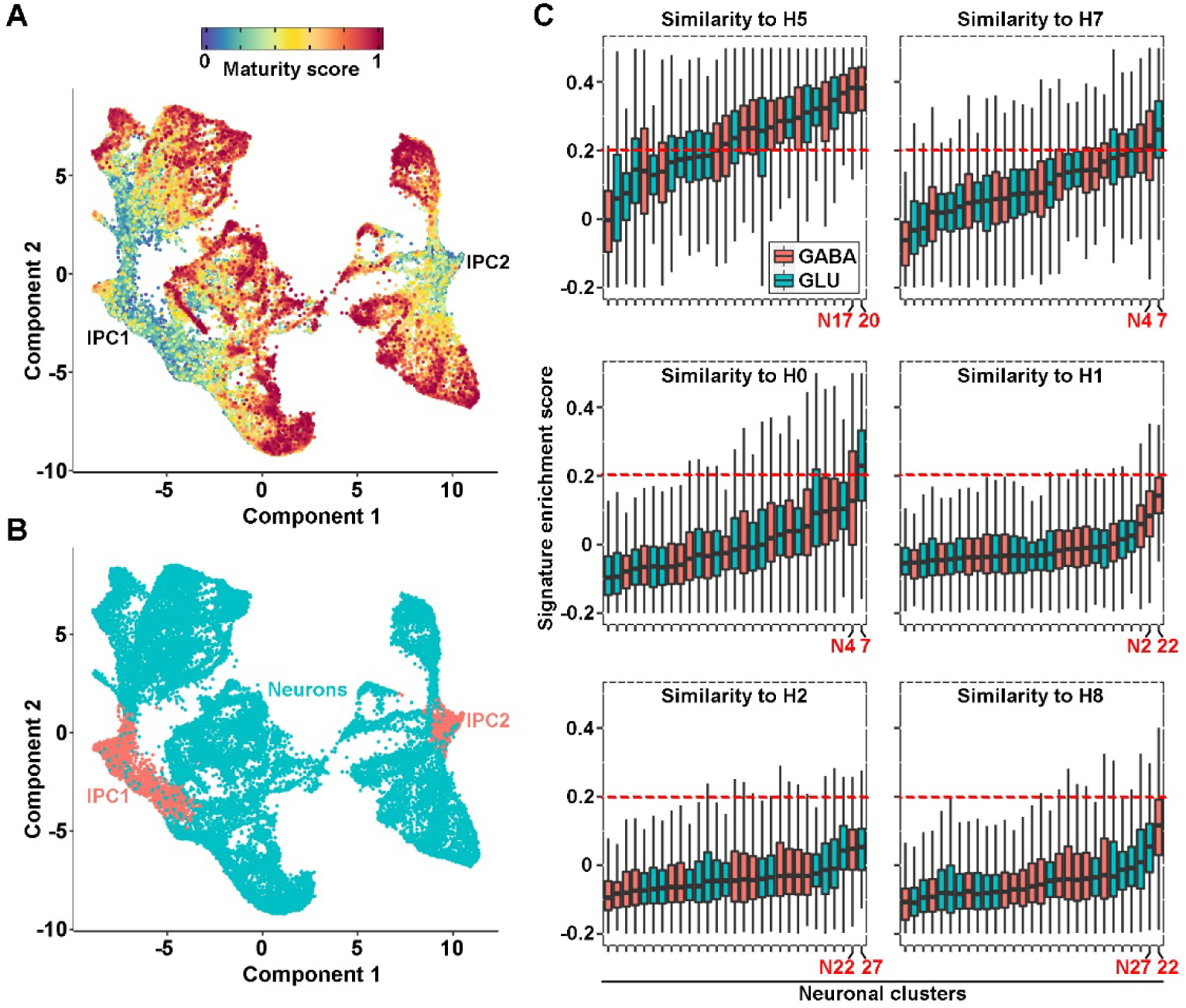
Progressive Differentiation from HIPCs, Nascent Neurons to Relatively Mature Neurons. Related to Figure 4. (A) Differentiation state of developing hypothalamic neurons in combination with IPCs that is predicted by CytoTRACE and overlaid on UMAP representation. (B) UMAP visualization of IPC subpopulations and neurons. (C) Enrichment of HIPC subtype (H0-2, H5, H7-8) signatures for each neuronal subtype. The signature transcriptional programs in each HIPC subtype were derived from the top 50 feature genes defining HIPC subtype identity (Table S6), and used to score the signature enrichment of 29 neuronal subtypes. The enrichment scores of H1, H2 and H8 signatures in all neuronal subtypes are very low, confirming the primitive state of these HIPC clusters.

**Figure S11.**
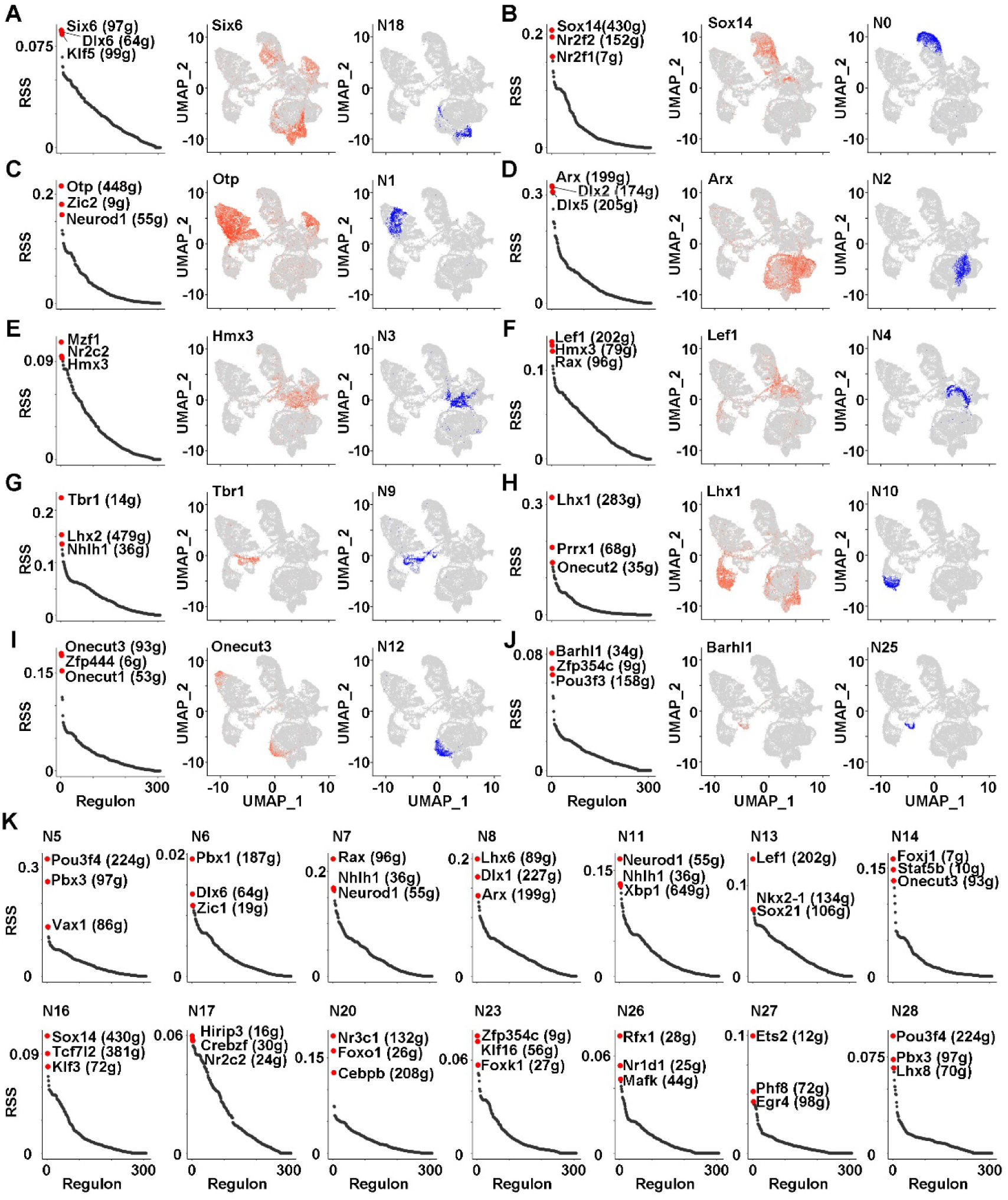
Neuronal Subtype-Specific Regulons. Related to Figure 5. (A-J) Shown are the ranked regulons based on regulon specificity score (RSS), UMAP projection of top regulon and UMAP representation of specific neuronal subtypes (from left to right). The top 3 regulons for each neuronal subtype are labeled and the number of predicted target genes are shown in the brackets. (K) Shown are the ordered regulons in diverse neuronal subtypes.

**Figure S12.**
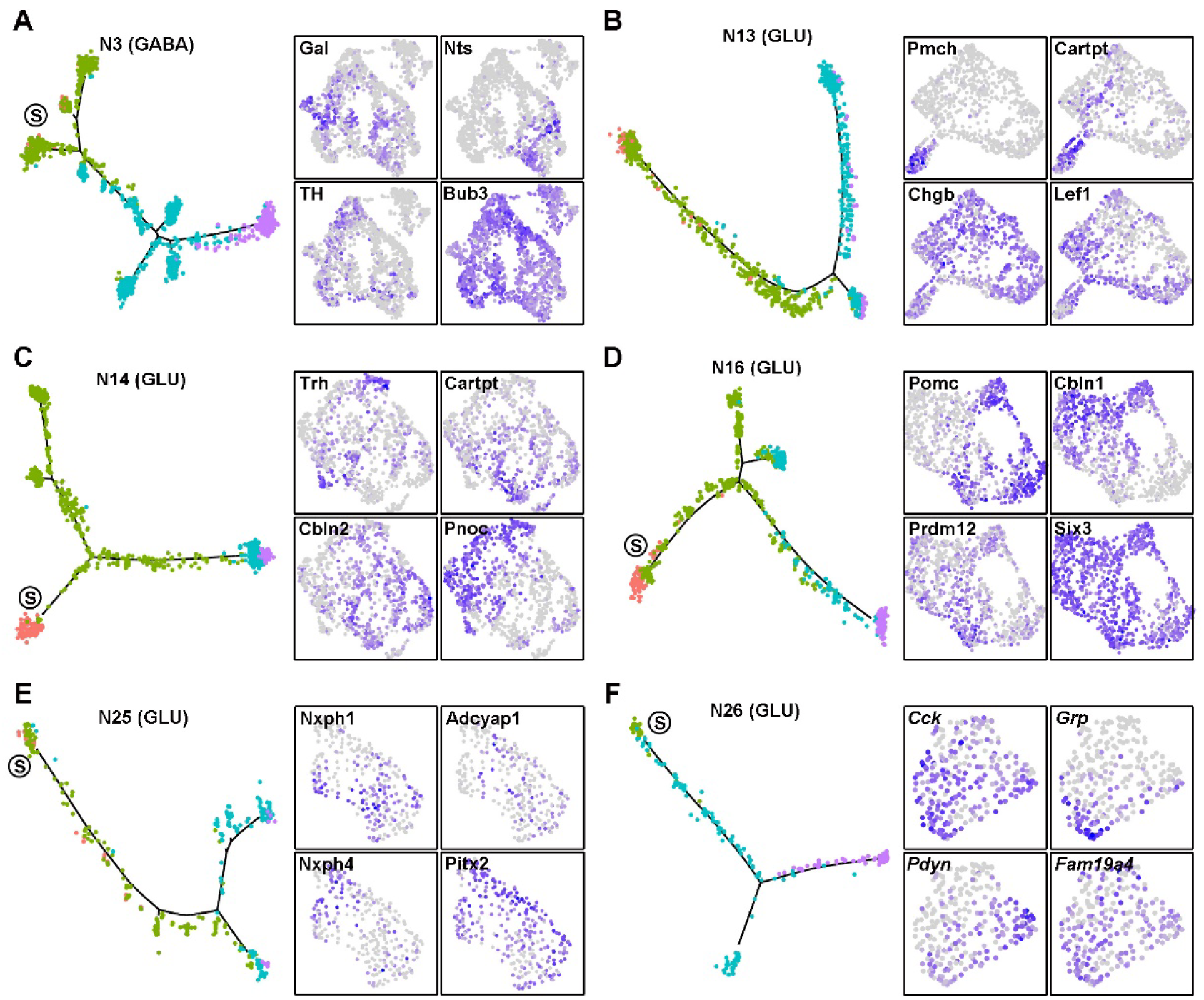
Fate Diversification of Hypothalamic Peptidergic Neurons During Maturation Process. Related to Figure 6. (A-F) Pseudotemporal trajectory of representative neuronal subtypes (left) and the UMAP projection of respective neuropeptide/TF expression (right).

**Supplementary Table 1. Top 50 Differentially Expressed Genes Enriched for Each Hypothalamic Cell Type (Related to Figure S2).**

**Supplementary Table 2. Top 100 Differentially Expressed Genes Enriched in E11, E14, P0 and P7 for Hypothalamic Cells (Related to Figure S2).**

**Supplementary Table 3. Top 100 Differentially Expressed Genes Enriched in E11, E14, P0 and P7 for RGCs or RGC-like cells (Related to Figure S3).**

**Supplementary Table 4. Top 50 Differentially Expressed Genes Enriched for Each RGCs Cell Type (Related to Figure S3).**

**Supplementary Table 5. Top 100 Differentially Expressed Genes Enriched in E11, E14, P0 and P7 for IPCs (Related to Figure S4).**

**Supplementary Table 6. Top 50 Differentially Expressed Genes Enriched for Each IPCs Cell Type (Related to Figure S5).**

**Supplementary Table 7. Top 50 Differentially Expressed Genes Enriched for Each Neuronal Subtype (Related to Figure 4).**

**Supplementary Table 8. Gene List of Mouse Transcription Factors (Related to Figure 5).**

**Supplementary Table 9. Gene List of Neuropeptides and Monoamines (Related to Figure 6).**

**Supplementary Table 10. The Sequences of HCR Probes Used in This Study.**

